# Applications of Boolean modeling to study the dynamics of a complex disease and therapeutics responses

**DOI:** 10.1101/2023.04.05.535790

**Authors:** Ahmed Abdelmonem Hemedan, Reinhard Schneider, Marek Ostaszewski

**Affiliations:** Luxembourg Centre for Systems Biomedicine, University of Luxembourg, Esch-sur-Alzette, Luxembourg

**Keywords:** Boolean modelling, Molecular mechanisms, Therapeutic target, Systems medicine

## Abstract

Computational modeling has emerged as a critical tool in investigating the complex molecular processes involved in biological systems and diseases. In this study, we apply Boolean modeling to uncover the molecular mechanisms underlying Parkinson’s disease (PD), one of the most prevalent neurodegenerative disorders. Our approach is based on the PD-map, a comprehensive molecular interaction diagram that captures the key mechanisms involved in the initiation and progression of PD. Using Boolean modeling, we aim to gain a deeper understanding of the disease dynamics, identify potential drug targets, and simulate the response to treatments. Our analysis demonstrates the effectiveness of this approach in uncovering the intricacies of PD. Our results confirm existing knowledge about the disease and provide valuable insights into the underlying mechanisms, ultimately suggesting potential targets for therapeutic intervention. Moreover, our approach allows us to parametrize the models based on omics data for further disease stratification. Our study highlight the value of computational modeling in advancing our understanding of complex biological systems and diseases, emphasizing the importance of continued research in this field. Furthermore, our findings have potential implications for the development of novel therapies for PD, which is a pressing public health concern. Overall, this study represents a significant step forward in the application of computational modeling to the investigation of neurodegenerative diseases, and underscores the power of interdisciplinary approaches in tackling challenging biomedical problems.

## 1 INTRODUCTION

The interpretation of omics data is crucial to develop meaningful hypotheses to understand complex disease mechanisms (Valle (2019)). To interpret these data, pathway databases play a role in providing an overview of the processes involved in disease mechanisms. Community-driven initiatives, such as disease maps, have been established to encode disease mechanisms in a computable form (Mazein (2018)). The disease maps can be further visualized through the use of dedicated biocuration and visualization tools (Funahashi (2007); Balci (2020); Wiese (2004); Kuperstein (2013); Gawron (2016)). The information provided by pathway databases and disease maps can then be utilized to develop computational models, which are crucial in advancing our understanding of the dynamic properties of complex diseases (Silk (2014)). By integrating experimental data with computational models, researchers can identify key molecular players and pathways involved in disease initiation and progression, and predict the effects of therapeutic interventions. Therefore, the use of disease maps and computational modeling represents a promising approach in the field of biomedical research, with potential implications for the development of personalized medicine and precision therapies Dynamic modeling approaches such as Boolean or Multi-valued models, Petri nets, and Ordinary Differential Equations (ODEs) (Dubrova (2006); Aalst (2009); Walter (1998)) can be used to represent the complex dynamics of biological systems and diseases. However, parameterizing these models remains a significant challenge (Ilea (2012)). Logical models provide a valuable alternative for researchers, as they can be more easily constructed and parameterized (Naldi (2018)).

By utilizing logical models, disease maps can be encoded in a computable form, and the dynamics of diseases can be analyzed in a more straightforward manner. These logical models can be further integrated with experimental data to create more comprehensive and accurate representations of disease dynamics. This approach has the potential to facilitate the discovery of novel drug targets and the development of personalized medicine.

Therefore, further research is needed to better understand the molecular mechanisms underlying diseases and advance personalized medicine. By leveraging community-driven initiatives such as disease maps and computational modeling approaches, researchers can gain a deeper understanding of complex diseases, and ultimately, develop more effective treatments.

Logical modeling in systems biology is a mathematical approach that uses Boolean algebra to represent the interactions between components in a biological system (Albert and Thakar (2014)). Logical modeling is a valuable tool for understanding the behavior of cellular networks and gene regulation networks, as well as for predicting the effects of perturbations such as drug treatments (Maldonado et al. (2017)). There are several types of logical modeling used in systems biology, including Boolean models (BMs), probabilistic BMs, Petri nets, and rule-based models (Aalst (2009)).

Boolean models are the most commonly used type of logical modeling in systems biology (Albert and Thakar (2014)). Boolean models represent variables with binary values of one (ON) or zero (OFF) (Albert and Thakar (2014)). The behavior of the output biomolecules is described by Boolean functions (BFs) that define the interactions between the inputs. Updating schemes determine the order in which BFs are calculated (Hemedan et al. (2022)). BFs have been primarily used to describe gene regulation, but they have also been applied to signaling networks using various logical formalisms, including Boolean, differential, and fuzzy equations (Terfve (2012); Bloomingdale (2018); Eduati (2020)).Boolean models provide a qualitative representation of the system and its interactions, without the need for detailed kinetic information (Aalst (2009)). However, in some cases, presenting data in a simple BM may not provide the best description of the biological system, and integration with other quantitative methods may be necessary (Maldonado et al. (2017)).

In this study, we utilized Boolean modeling to represent the complex molecular mechanisms underlying Parkinson’s disease (PD). By using the Parkinson’s disease map Fujita et al. (2014) PD mechanisms in a computable form, we simulated the dynamics of the disease and proposed potential drug targets. By abstracting the disease mechanisms in a logical form, we can simulate disease dynamics and identify potential therapeutic targets, ultimately facilitating the development of personalized medicine. For example, LRRK2 mutations have been found to increase the aggregation of cytosolic proteins, leading to apoptosis and cell dysfunction, which could be targeted by therapeutic interventions (Gopalakrishna and Alexander (2015)).

The PD map is translated into BMs in an automated fashion using CaSQ tool (Aghamiri et al. (2020)). The complexity of disease pathways is studied by simulating the effect of genomic burden from omics data. BMs are created at different scales of the complexities to ensure its ability and reliability to simulate disease mechanisms. These different levels of complexity refer to the different scales at which disease pathways can be represented and analyzed. First, the simple and known mechanisms are simulated to investigate the model ability to represent the already known behaviour. Further, the BMs are used to re-simulate complex molecular interventions data, comparing the results with the literature. Creating and validating BMs at different levels of complexity is a crucial step towards developing accurate simulations of biological processes. Gradually increasing the complexity of the model allows us to identify key factors that impact the accuracy of the simulation, such as specific biological components and interactions, and refine the model accordingly. Through this process, we can better understand the complexity of disease mechanisms and develop more reliable results that can ultimately contribute to the development of effective treatments. BMs were created in different modelling formats, facilitating the interoperability and the simulation across different tools. This includes SBML-qual, which is used for creating, storing and exchanging qualitative models (Chaouiya et al. (2013)). Analysis of the models’ structural and dynamic properties was used to verify their accuracy. The dynamic verification and sensitivity analysis results showed that the BFs accurately represented the original interactions and the model was robust against small perturbations. The validation was performed by simulating models in different scales of complexity. We analyzed and validated a number of pathways based on the literature and experimental evidence. These included the TCA cycle, dopamine transcription, FOXO3, and the Wnt-PI3k/AKT signaling pathway. Our results demonstrated the utility of Boolean modeling in simulating the behavior of these pathways and predicting the effects of perturbations, providing valuable insights into the molecular mechanisms underlying PD.

## 2 MATERIAL AND METHODS

### 2.1 Construction of a Boolean model

The diagrams analyzed in this paper were obtained from the MINERVA Platform (Gawron (2016)) (see the Supplementary Materials). This platform provides the capability to export specific diagrams from the map. The diagrams in CellDesigner SBML format were then automatically transformed into SBML-qual format using the CaSQ (CellDesigner as SBML-qual) tool. The reduction of the diagrams was performed in accordance with the diagram reduction rules outlined in Aghamiri et al. (2020).

The translation process involves converting the Process Description notation into the Activity Flow notation (see fig. 1). Biomolecules in Process Description diagrams may exist in various states (such as phosphorylated or methylated), however, during the translation to the Activity Flow notation, each biomolecule is represented as a single node with its different states depicted as distinct logical states of the node, such as active or inactive.

**Figure 1.**
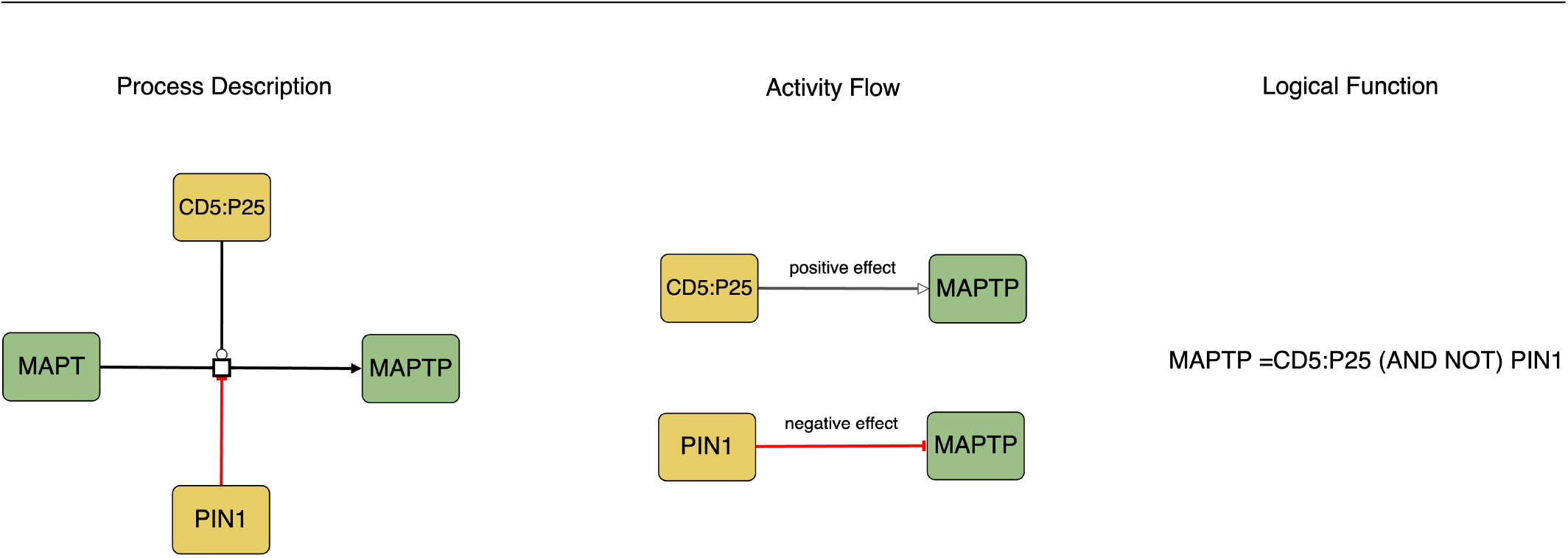
This figure represents the causal molecular interactions in both Process Description and Activity Flow. The logic equation depicted in the Process Description indicates that the activity of MAPTP as a product is determined by the presence of CD5:P25 and the absence of PIN1.

### 2.2 Topological analysis of the models

The structural and functional correctness of BM was evaluated by analyzing the interactions between the biomolecules. To this end the topological features of the BM were analyzed as a network. To do so, Simple Interaction Format (SIF) was created as a simple way to represent pairwise interactions among biomolecules in a network graph. To transform an SBML model into SIF format, the model is first converted into a graph representation with nodes representing biomolecules and edges representing interactions between biomolecules. Relevant information such as the source and target nodes of each edge is then extracted from the graph representation and mapped to the SIF format. The resulting SIF representation is a list of pairwise interactions between nodes, which can be useful for evaluating the structural correctness of biological models. Established tools Trinh and Kwon (2019), were employed to analyze the network, including the in/out degree, feedback loops, and centrality measures. Among the centrality measures, betweenness and stress centrality were utilized to quantify the significance of each node in the network.

Betweenness centrality measures the importance of a node as an intermediary between other nodes, by quantifying how frequently a node lies on the shortest path between pairs of other nodes in the network. Stress centrality measures the importance of a node in connecting other nodes by quantifying the number of shortest paths that pass through the node. (Ma’ayan (2011); Ashtiani et al. (2018)).

Betweenness centrality:

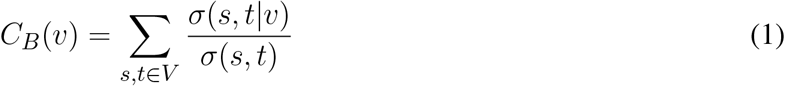

where *V* is the set of nodes in the network, *σ*(*s, t*) is the total number of shortest paths from node *s* to node *t*, and *σ*(*s, t*|*v*) is the number of those shortest paths that pass through node *v*.

Stress centrality:

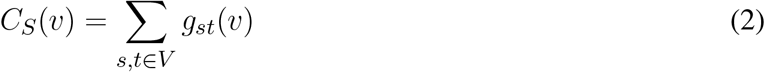

where *V* is the set of nodes in the network, and *g*_*st*_(*v*) is the number of shortest paths from node *s* to node *t* that pass through node *v*. Molecular targets for perturbation were prioritised based on their centrality and sensitivity measures. First, we computed betweenness centrality and stress centrality scores for all nodes in the network and identified the top-ranked nodes with high centrality values. Next, we simulated perturbations (Knockouts and overexpressions) to the network and measured the similarity and identity based distances between the original and perturbed states to assess the sensitivity of each node to the perturbations. By combining the centrality and sensitivity scores, we prioritized the molecular targets for perturbation that were highly central and sensitive.

### 2.3 Model analysis

#### 2.3.1 Model updating schemes

The updating schemes of Boolean functions (BFs) were evaluated and compared, including synchronous updating, asynchronous updating (Garg et al. (2008).

The synchronous updating scheme updates all biomolecules simultaneously according to their BFs (Garg et al. (2008); Wang (2012)

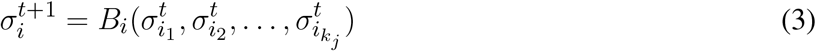

The asynchronous scheme updates the variables in a non-synchronous manner. The new value of each variable *σ* is denoted by an asterisk and is given by:

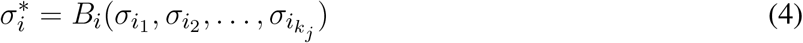

Here, the values of the inputs on the right-hand side of the equation can be either current or prior, depending on the individual timescales (Wang (2012)).

Probabilistic Boolean models (PBMs) was also be used to update BFs in a probabilistic manner by assigning probabilities to BFs and updating biomolecules based on these probabilities (Schwab (2020); Grieb (2015)). A PBM can be described as follows:

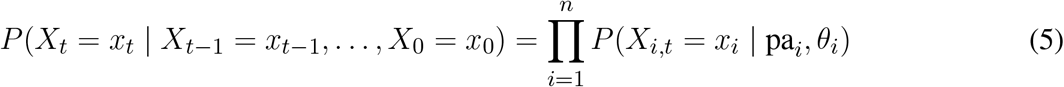

where *X*_*t*_ is the state of the network at time *t, x*_*t*_ is a vector of binary values representing the state of each node in the network at time *t*, pa_*i*_ is the set of parent nodes of node *i* in the network, and *θ*_*i*_ is a vector of probabilities representing the conditional probability distribution of node *i* given its parent nodes. The equation states that the probability of the network being in state *x*_*t*_ at time *t*, given its previous states and the network structure, is the product of the conditional probabilities of each node in the network.

Synchronous updating scheme was simulated using BoolNet tool in which all the biomolecules in the network updated simultaneously, based on the states of their regulators (Müssel (2010)). Asynchronous updating scheme was simulated using pyMaBoSS Stoll et al. (2017), see Stochastic Boolean model simulation section.

The behavior of a BM under different update schemes was visualized through state transition graphs (see section 5 in the supplementary file), which represent all possible states of the system and the transitions between them (Garg et al. (2008)). The state transition graph illustrates the range of outcomes for a given initial condition based on the update scheme used. Both update schemes demonstrated the ability to simulate expected system behavior.

#### 2.3.2 Attractor Search

Attractors are states in a state transition graph that have no outgoing edges and can be stable or complex. The basin of attraction is the set of states that lead to an attractor and can provide insight into potential biological scenarios (Hopfensitz (2012); Klemm and Bornholdt (2005)). During the verification of models, several search algorithms were examined to compare their speed and reachability. One method used was an exhaustive search, which aimed to find all possible attractors by performing synchronous transitions between states. To increase efficiency, a SAT-based technique was employed, where the problem of attractor identification was framed as a Boolean satisfiability problem (SAT). This allowed for determining if a particular formula was satisfiable or not and to limit the search to loops of a specific length (Biere (2008)). The SAT-based technique involves representing the state of the network as Boolean variables, and representing the Boolean rules that govern the transitions between states as logical constraints. Finding solutions to the SAT problem that correspond to attractors in the network allows for the attractor search.

The SAT-based technique can be formulated mathematically as follows:

Let *x*_1_, *x*_2_, …, *x*_*n*_ be the Boolean variables that represent the state of the network, where *x*_*i*_ is either 0 or 1. Let *f*_1_, *f*_2_, …, *f*_*m*_ be the Boolean functions that represent the transitions between states, where each function is a logical constraint that depends on the values of the Boolean variables. The goal of the SAT-based technique is to find a set of values for the Boolean variables that satisfy the logical constraints and correspond to an attractor in the network. This can be represented as a SAT problem as follows:

Find *x*_1_, *x*_2_, …, *x*_*n*_ such that 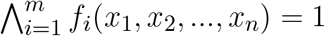

The SAT problem can be efficiently solved using SAT solvers to determine the attractors in the network.

Another approach, the decomposition method, aimed to optimize speed and reduce complexity by breaking down the model into strongly connected components (SCCs). The method can be represented mathematically as:

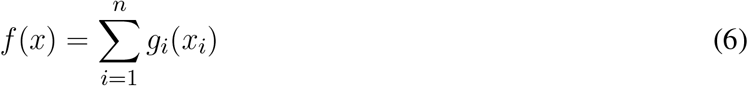

where *f* (*x*) is the original problem, *n* is the number of subproblems, and *g*_*i*_(*x*_*i*_) is the solution to each subproblem. The subproblems are usually defined so that they can be solved independently and then combined to obtain a solution to the original problem.

In addition, a heuristic search and an asynchronous search were conducted. The heuristic search involved selecting a subset of possible states as initial conditions and performing synchronous transitions until an attractor was reached. The asynchronous search used random asynchronous transitions to identify steady states and complex attractors from the chosen initial conditions.

#### 2.3.3 Perturbation analysis

) A perturbation analysis was conducted to evaluate the effect on the topological robustness, dynamic resilience, and attractors reached by the models (Trinh and Kwon (2019)). Specially, we focused on node perturbations, which altering the state of a single biomolecule through knockout and overexpressions.

The evaluation was performed by performing sensitivity analysis on a selected set of models, examining each biomolecule. Sensitivity analysis is a technique that assesses how changes in a model or system’s inputs affect its output, in this case, the two attractors reached by the model (unperturbed and perturbed). To quantify the difference between the two attractors, similarity-based distance and identity-based distance were used. Similarity-based distance assessed the similarity between the two attractors by taking into account the common and unique states in both. The similarity-based distance between the unperturbed attractor (*A*_*unpert*_) and the perturbed attractor (*A*_*pert*_) can be defined as:

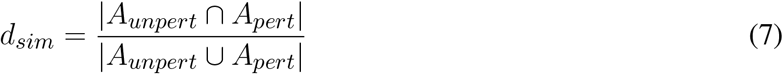

Identity-based distance, on the other hand, measured the percentage of states present in both attractors.

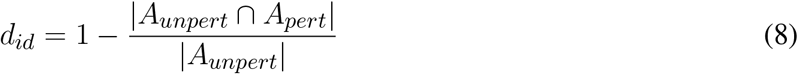

#### 2.3.4 Stochastic Boolean model simulation

The simulations of the specified biological models were conducted using pyMaBoSS, a python API for the MaBoSS software (Stoll et al. (2017)). This framework provides a tool for probabilistic Boolean modeling and simulation of biological systems through discrete/continuous time Markov processes. It operates by utilizing a Monte Carlo algorithm that simulates the system’s evolution over time based on the initial conditions of the biomolecules and the interactions between them.

pyMaBoSS performs asynchronous updates through a random walk simulation technique, in which a single biomolecule is selected and updated at each step, followed by repeating this process to obtain a sample of the attractors. Asynchronous transitions are used in pyMaBOSS to determine steady states and complex attractors from the specified initial states.

The probabilities of the initial states are calculated based on the effect size and statistical correlations of the PPMI dataset, which are used to assign probabilities to the initial states. These probabilities are then included in configuration files for use with PyMaBoSS.

In pyMaBoSS, a biological system is represented by a model of interconnected Boolean variables, with each variable representing the state of a biomolecule (e.g. present or absent, active or inactive). The interactions between the variables are defined by Boolean rules, which indicate the impact of one variable’s state on another. The Monte Carlo algorithm randomly samples the possible states of the system at each time step, based on the current state of the system, the probabilities of each state, and the interactions between the variables. By simulating the system over multiple time steps, pyMaBoSS estimates the probability of each state at each time point (Stoll et al. (2017)).

## 3 RESULTS

We investigated the dynamic behavior of molecular mechanisms in Parkinson’s disease (PD) and their response to changes. To achieve this goal, a variety of BM formats - including SBML-qual - were created and analyzed using a range of tools. The validity of the BMs was evaluated by examining their structural and dynamic properties. The results of this assessment demonstrated that the models displayed consistent structural and dynamic properties, suggesting their use in study of the dynamics of PD.

### 3.1 Model construction

To construct the Boolean models of PD mechanisms, we first selected a group of PD map diagrams for downstream modeling and verification. We carefully chose diagrams that were directly relevant to PD-related phenotypes, as outlined in table 1 (see also figures S1 to S6 i n the supplementary file). Once we identified the relevant pathways from the PD map, we exported them in CellDesigner SBML format, allowing for further analysis and modeling. The CellDesigner SBML diagrams provided us with a detailed representation of the biological pathways involved in PD, which we used to construct our Boolean models.

**Table 1.** This table presents a summary of the selected pathways with their nodes, and edges in several important cellular signaling and metabolic pathways. The pathways listed include dopamine transcription, Wnt-PI3K/AKT signaling, FOXO3 activity, mTOR-MAPK signaling, PRKN mitophagy, PPARGC1A, and the TCA cycle. The number of nodes and edges for each pathway is also provided

| Pathways | Nodes | Edges | Type |
| --- | --- | --- | --- |
| Dopamine transcription | 167 | 196 | Cellular signalling |
| Wnt-PI3K/AKT signalling | 391 | 436 | Cellular signalling |
| FOXO3 activity | 65 | 86 | Cellular signalling |
| mTOR-MAPK signalling | 59 | 83 | Cellular signalling |
| PRKN mitophagy | 54 | 72 | Cellular signalling |
| PPARGC1A | 67 | 109 | Cellular signalling |
| TCA cycle | 137 | 160 | Metabolic |

To generate the SBML-qual models necessary for our modeling work, we used CaSQ. This software allowed us to convert the CellDesigner SBML diagrams into a SBML-qual model that facilitated the interoperability between modeling tools. With these Boolean models, we analysed the PD mechanisms to explore the underlying biology of the disease and gain insights into potential therapeutic targets.

### 3.2 Model verification

To verify the accuracy and reliability of BMs, both structural and dynamic aspects of the model must be examined. First, we evaluated the interactions and relationships between the model’s components, to examine how they reflect the underlying biological system. For dynamic verification, we analysed the model’s behavior over time and compared it with the known biological behaviour represented in the literature.

#### 3.2.1 Sensitivity and structural analysis

We aimed to identify molecules that can serve as potential targets and intervention points in pathways. To achieve this, we conducted a structural and sensitivity analysis. We selected the molecules with the highest betweenness centralities of each respective model (top quantile). These molecules were proposed to play a role in maintaining the stability of the system as they are located at the intersection of multiple paths. The table below (table 2) shows the identified molecules and their corresponding betweenness centralities.

**Table 2.** The table displays the betweenness centrality values and knockout sensitivity for key pathways and relevant biomolecules. The bolded valued indicate that biomolecules with high betweenness have low knockout sensitivity, and vice versa, while some with low betweenness have high knockout sensitivity

| Pathway | ID | Betweenness centrality | Knockout sensitivity |
| --- | --- | --- | --- |
| PPARGC1 | PPARGC1A phosphorylated | 322 | **0.14** |
|  | TF NRF1 complex | 174 | 0.04 |
|  | PPARGC1A acetylated phosphorylated | 94 | 0.29 |
|  | TF NRF2 complex | 88 | 0.11 |
|  | TF YY1 complex | 75 | **0.58** |
| Wnt/Pi3kAKT | mTORC1 complex neuron | 463 | **0.03** |
|  | AKT1 phosphorylated | 440 | 0.04 |
|  | PDPK1 | 260 | 0.09 |
|  | RPS6KB1 phosphorylated | 189 | 0.07 |
|  | IRS1 phosphorylated | 173 | **0.43** |
| Mtor | AMPK complex neuron | 2276 | **0.42** |
|  | TSC1:TSC2 complex neuron | 1465 | 0.60 |
|  | STK11 | 1018 | 0.47 |
|  | SESN2 | 788 | 0.60 |
|  | nicotinamid | 745 | **0.50** |
| FOXO3 activity | TF space FOXO3 complex nucleus | 3673 | 0.49 |
|  | TF space CHOP:FOXO complex | 1762 | 0.47 |
|  | FOXO3 nucleus | 1600 | **0.46** |
|  | SIRT3 | 1482 | 0.47 |
|  | BBC3 rna | 895 | **0.59** |
|  | BCL2L11 rna | 895 | 0.42 |
| Dopamine transcription | TF NR4A2 complex | 329 | **0.17** |
|  | PITX3 | 101 | 0.16 |
|  | TF PITX3 complex | 66 | 0.14 |
|  | MIR133B rna | 54 | 0.17 |
|  | RXRA | 26 | **0.50** |

The second step of our analysis aimed to evaluate the sensitivity of the identified high central molecules to perturbations, specifically gene knockouts. We investigated whether these molecules are reliable measures to estimate the effect of perturbation and to identify potential targets for intervention.

We found that most of the high central molecules showed less sensitivity against knockouts, indicating their robustness in maintaining the system’s stability. This observation suggests that targeting these molecules may not have a significant impact on the overall system stability. However, this does not necessarily mean that they are not potential targets for intervention, as their perturbation may have downstream effects that affect other pathways or processes (see fig. 2).

**Figure 2.**
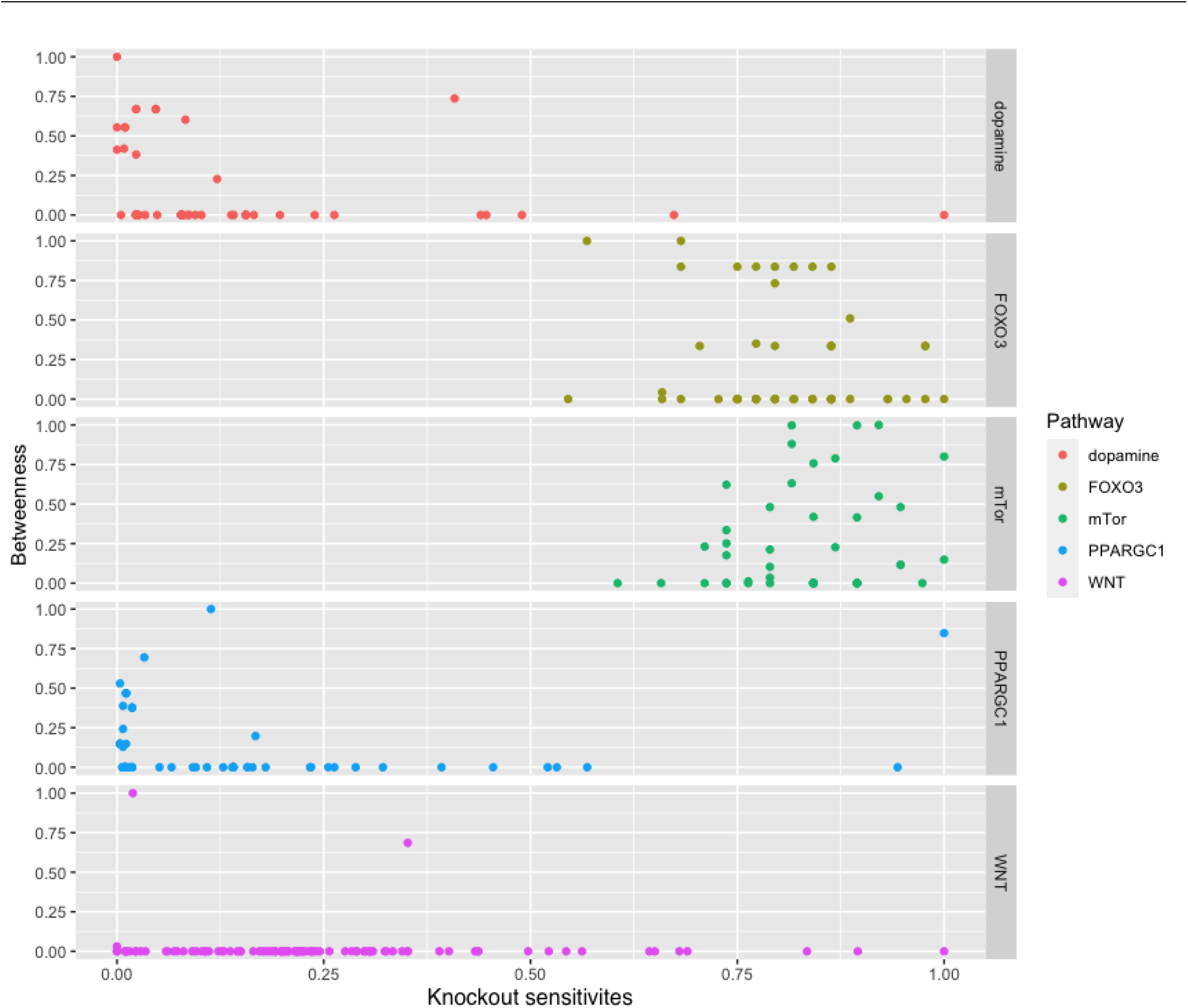
The figure displays a comparison between the betweenness centralities and sensitivity measures in selected pathways. While most pathways exhibit high betweenness centralities, their sensitivity measures are low. This suggests that betweenness centrality may not be a reliable indicator of the significance of biomolecules in these pathways.

To further investigate the reliability of the identified molecules as potential targets for intervention, we identified the common molecules that have significant centralities and/or sensitivities and also have reported biological importance (see tables 3 and 4. These perturbations were selected based on known biological scenarios to assess the model’s ability to reproduce the known behavior.

**Table 3.**
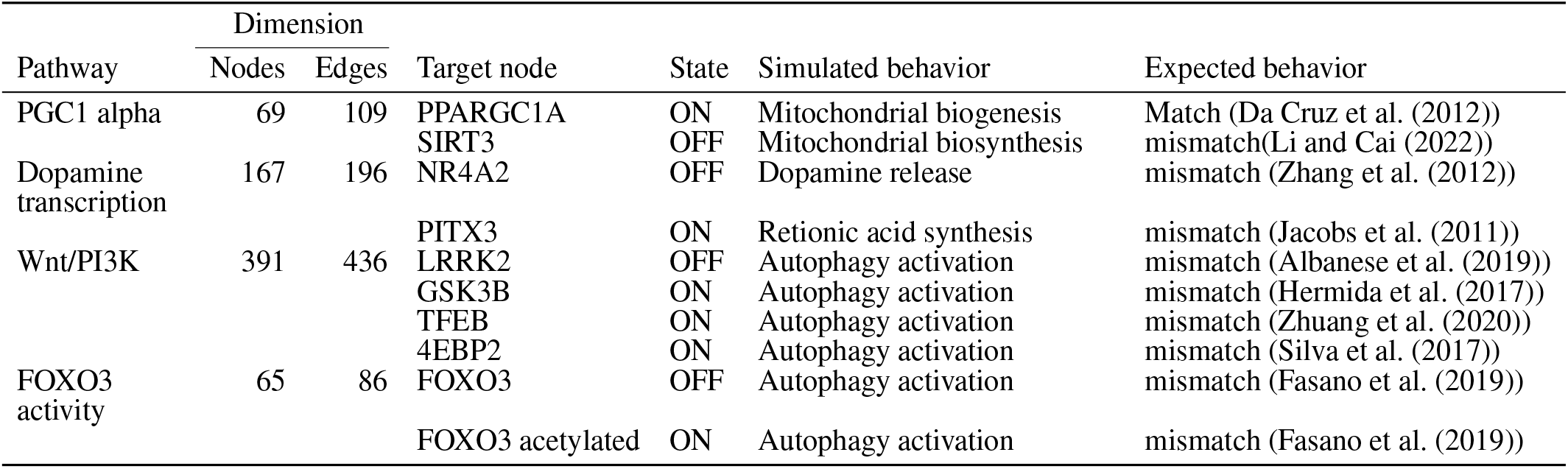
The table compares the simulated behavior of decomposed Boolean models to expected behavior based on published literature. The table includes information on the pathways, the number of nodes and edges in each network, the target node, the state of the target node (ON or OFF), and the simulated and expected behavior for each pathway.

**Table 4.**
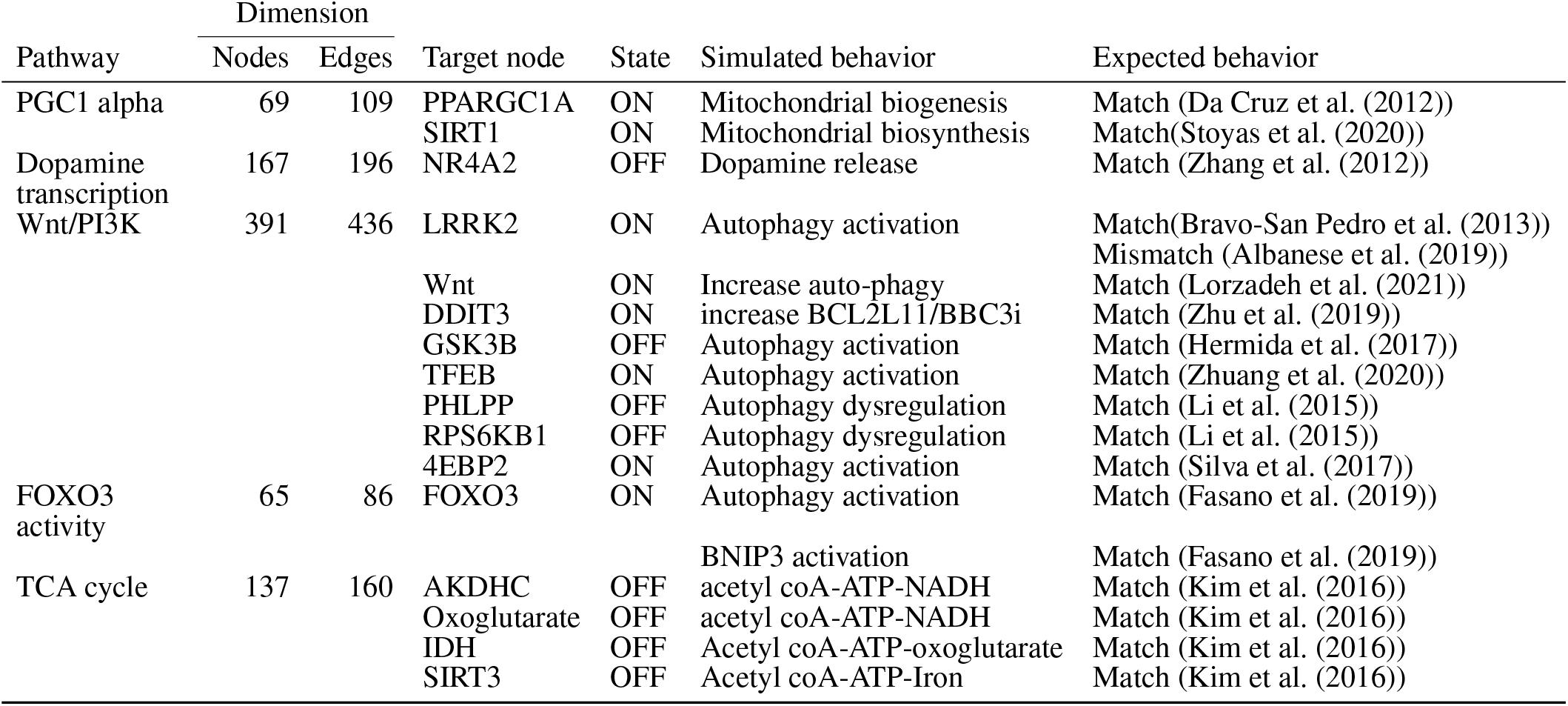
The table compares the simulated behavior of several Boolean models to expected behavior based on published literature. The table includes information on the pathways, the number of nodes and edges in each network, the target node, the state of the target node (ON or OFF), and the simulated and expected behavior for each pathway.

In the Wnt Pi3kAKT pathway, we identified molecules such as ROCK2, EIF4E, IGFR1, IGF1, and INS that can compensate for the absence of PDPK1, RPS6KB1 phosphorylated (fig. 3). The presence of such alternative molecules can maintain the overall function of the pathway when such a scenario arises. These compensatory paths may be exploited to develop targeted interventions for pathological conditions.

**Figure 3.**
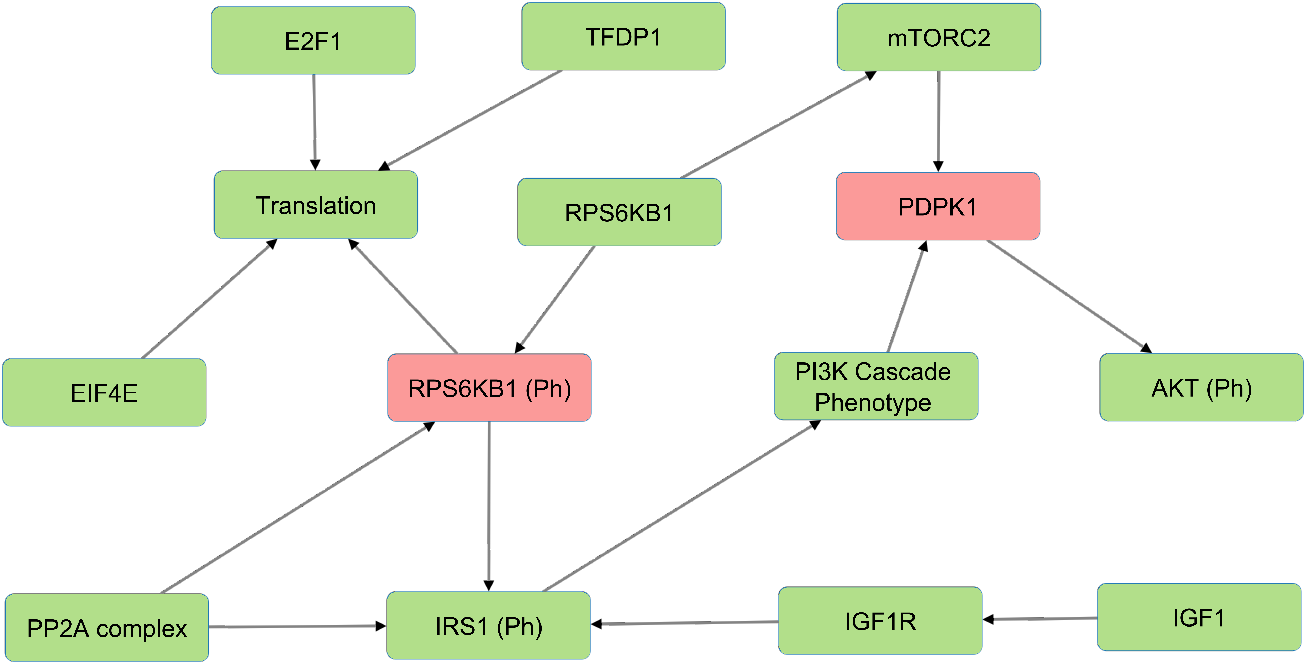
The figure illustrates an example of alternative molecules in the Wnt/PI3K-AKT model (shown in green) that compensate for PDPK1 and RPS6KB1 phosphorylated knockouts (shown in red) and reduce sensitivities. These compensatory molecules enable the pathway to continue functioning despite the loss of the phosphorylated proteins and reduce the overall sensitivity of the pathway to perturbations.

#### 3.2.2 Attractor reachability

In this study, we aimed to identify stable states, known as attractors, within the analyzed pathways using Boolean modeling. To ensure the biological relevance of the identified attractors, a thorough literature search was conducted. The search aimed to identify experimental evidence for the molecular states and interactions observed in the identified attractors. The findings of this literature search were summarized in tables 3 and 4, which present the expected behavior of the molecules and their corresponding stable states. The relevance of each stable state was evaluated based on its correspondence to known biological states, its association with PD, and its relevance to the analyzed pathways.

Our results indicated that the attractors produced by decomposition-based approaches (in methods) were not biologically relevant for all pathways (see fig. 4 and table 3). This discrepancy was attributed to their aggressive decomposition which resulted in overly fragmented models (available in the Gitlab repository).

**Figure 4.**
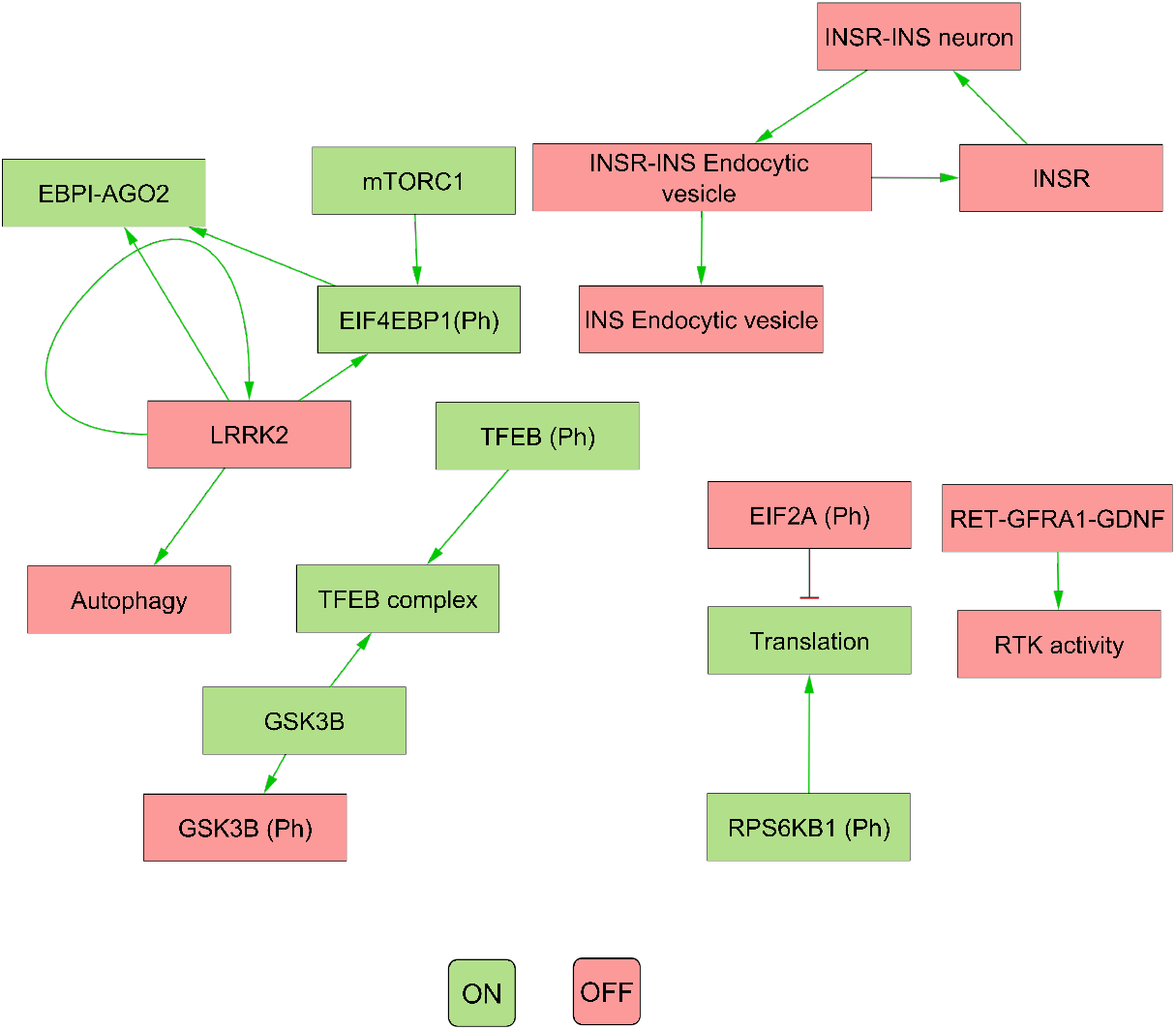
The figure illustrates the attractor pattern of decomposed WNT-Pi3k/AKT. The yellow and green colours represent the OFF and ON states of the molecules in the attractor

However, the attractors produced by Heuristic and SAT solver algorithms were found to be viable for all pathways except for the Wnt/PI3K pathway (see table 4). Therefore, we deemed Heuristic and SAT solver algorithms more suitable for further analysis.

### 3.3 Validation of Boolean models from literature

We validated the model outputs by comparing them to the literature. We first selected targets for the pathways and perturbed them according to scenarios suggested by literature. We then simulated the changes in the corresponding outputs. This was achieved through: i) running a simulation with modeling tools and the CellCollective platform Helikar (2012); Trinh and Kwon (2019); Stoll et al. (2017). ii) Comparing the computed attractors by modeling tools with the differential expressed molecules reported in corresponding published dataset (Zhang et al. (2012)) and literature as shown in table 4. The validation process has yielded both matching and mismatched scenarios between the model outputs and pertinent literature. Further exploration into the causes of the observed mismatches will contribute to an enhanced understanding of the limitations and strengths of the model. Further, the comparison to literature, as described, represents a crucial step in the ongoing refinement of our comprehension of the pathways and perturbations. The validation scenarios, results, and interpretation for the selected pathways are discussed in subsequent sections.

#### 3.3.1 Validation of TCA cycle model with literature and experimental evidence

The TCA cycle, a metabolic pathway occurring in the mitochondria of cells, is disrupted in Parkinson’s disease (PD), potentially contributing to its development (Shen et al. (2020)). This pathway is an ideal candidate for validating the BM approach due to the extensive literature and data available for comparison, as well as its well-studied and understood nature (Ahn et al. (2017)). Furthermore, the TCA cycle involves the transformation of multiple metabolites, which can be monitored through techniques such as fluxomics, providing a means of evaluating the BM’s accuracy by comparing predicted and experimentally observed metabolite levels. Given its central role in various cellular processes, a BM of the TCA cycle could have broad and far-reaching implications.

The validity and precision of the TCA cycle BM are established through a comparison with literature and fluxomics data. To run the simulation, the knockouts of isocitrate dehydrogenase, alpha-ketoglutarate dehydrogenase, and the pyruvate dehydrogenase complex are selected as the initial state parameters based on prior research (Ahn et al. (2017)). Using temporal-fluxomics data, the levels of alpha-ketoglutarate dehydrogenase activity and the nucleotide triphosphates GTP and GDP’s impact on ATP levels are modeled using CellCollective. The simulation results are then compared to experimental data as described below.

As shown in the fluxomics data, the validated results based on literature indicate that the simulation of increased activation of acetyl CoA, NADH, and PDKs leads to an increase in phosphorylation reaction, subsequently reducing the activity of the pyruvate dehydrogenase complex (PDC). This finding supports the hypothesis that PDC deficiency, a potential target for therapeutic intervention in age-related diseases, arises from the activation of the phosphorylation reaction involving these substances (Stacpoole (2012)). The deficiency of PDC, caused by KGDHC knockout, has a significant impact on the production of succinic semialdehyde (Sergi and Parayil Sankaran (2021)). This deficiency results in decreased levels of succinic acid and succinyl CoA, leading to a decline in ATP and GTP in mitochondria (Shi et al. (2011); Gibson et al. (2003)). The knockout of isocitrate dehydrogenase leads to a decrease in ATP levels and a disruption in the oxidative decarboxylation catalysis of isocitrate into alpha ketoglutarate, resulting in mitochondrial dysfunction and dopaminergic neurotoxicity (Kim et al. (2016)). In PD, key regulators such as oxoglutaric acid, glutamate hydrogenase 1 (GLUD), and ATP levels are disturbed in response to SIRT3 knockout, which directly impacts mitochondrial function (Shen et al. (2020)). These findings are summarized in table 4.

The cellular metabolites undergo fluctuations during the cell cycle, adapting to changes within the cell. The impact of alpha KGDH, GTP, and GDP on ATP levels are modeled in the simulation. The simulated ATP activity levels match the measured concentrations in synchronized HeLa cells every 3 hours following release from growth arrest

#### 3.3.2 Dopamine transcription pathway

One of the pathways validated in this study was the dopamine transcription pathway, known to be disrupted in Parkinson’s disease (PD) (Barneda-Zahonero et al. (2012)).

NR4A2 were selected as the key elements in this validation, as they are proteins that play crucial roles in the development and maintenance of neurotransmitters and various cellular processes (Barneda-Zahonero et al. (2012); Li et al. (2020)).

In the simulation, the effects of perturbations to NR4A2 and SIRT1 on dopamine release and mitochondrial biosynthesis were observed as activity levels using CellCollective. These results, confirmed through a comprehensive attractor analysis (available in the Gitlab repository), demonstrate the following behavior in line with literature (table 4):

- The production of brain-derived neurotrophic factor (BDNF), crucial for the survival and growth of neurons, was impacted by the NR4A2 knockout. This result aligns with previous research showing that NR4A2 is involved in the stimulation of BDNF production in response to the neurotransmitter N-methyl-D-aspartate (NMDA) (Barneda-Zahonero et al. (2012)).
- The NR4A2 knockout was also observed to affect other molecules such as GCH1, TH, DDC, SLC18A2, SLC6A3, and DRD2, which play significant roles in the development and maintenance of neurotransmitters through various targets, as previously reported in Jankovic et al. (2005); Kadkhodaei et al. (2013); Jacobs et al. (2009).

#### 3.3.3 Wnt-PI3K/AKT

The Wnt-PI3K/AKT pathway was evaluated in this study as it has been reported to be impacted by mutations in Parkinson’s disease (PD) (Madureira et al. (2020); Bravo-San Pedro et al. (2013); Rabanal-Ruiz et al. (2017)). The key elements of the pathway considered during the validation were LRRK2 G2019S mutation, Wnt, DDIT3, GSK3B, TFEB, PHLPP, RPS6KB1, and 4EBP2, which are involved in the development and progression of PD (Bravo-San Pedro et al. (2013); Poret and Guziolowski (2018); McCabe et al. (2020); Zhu et al. (2019)).

The simulation scenario involved perturbing these biomolecules and observing the effects on autophagy and neuron survival as activity levels using CellCollective. These results were confirmed through an exhaustive attractor search analysis. The results showed the following behaviors that are in line with the published literature (table 4, (table 6)):

- The simulation results showed that overexpression of the LRRK2 G2019S mutant increased autophagy (Bravo-San Pedro et al. (2013)).
- This increase was found to be mediated by the inhibition of mTORC1/2, which are proteins that regulate autophagy (Poret and Guziolowski (2018); McCabe et al. (2020)). These findings suggest that the interconnectedness of amino acid sensing, mTORC1 signaling, and autophagy may offer a promising approach for treating PD (Rabanal-Ruiz et al. (2017)).
- Activating both 4EBP2 and TFEB was found to increase autophagy activity more than activating them separately (Franco-Juárez et al. (2022); Decressac et al. (2013); Moors et al. (2017)). Specifically, activating both of these proteins in combination resulted in a 10.7% and 13.6% increase in autophagy compared to activating them individually.
- In addition, the combination of inhibiting the proteins RPS6KB1 and PHLPP and activating TFEB significantly decreased neuronal death and the active state of autophagy (Li et al. (2015)). This combination resulted in a 96.3% decrease in neuronal death.
- Both activating the Wnt protein and inhibiting the protein GSK3B were found to increase autophagy (Palomer et al. (2019); Hermida et al. (2017); Yang et al. (2018)). The combination of these modulations could improve our understanding of therapeutic protocols for neurological diseases by promoting neurogenesis and autophagy (Siegle (2018)). These findings suggest that targeting these proteins could be a promising approach for developing treatments for neurological disorders.

**Table 5.** This table presents data on the ATP activity in the TCA (tricarboxylic acid) cycle with fluxomics integration. The data includes the time in hours, the condition (labeled as “Cond.”), and the levels of various inputs (alpha-ketoglutaric acid, GTP, GDP, and phosphate). The table also includes both simulated and real measurements of ATP levels.

| Time (h) | Cond. | Inputs |  |  |  | Output ATP level |  |
| --- | --- | --- | --- | --- | --- | --- | --- |
| | | α-KG acid | GTP | GDP | Phosphate | Simulated | Real |
| 9 | C1 | 0.84 | 0.48 | 0.50 | 0.45 | 1 | 0.87 |
| 12 | C2 | 0.79 | 0.77 | 0.66 | 0.67 | 0.86 | 0.77 |
| 15 | C3 | 0.77 | 0.98 | 0.79 | 0.93 | 1 | 1 |
| 18 | C4 | 0.84 | 0.83 | 0.83 | 0.71 | 1 | 0.95 |
| 21 | C5 | 1 | 0.56 | 0.65 | 0.73 | 1 | 0.90 |
| 24 | C6 | 0.88 | 0.50 | 0.64 | 0.72 | 0.96 | 0.84 |
| 27 | C7 | 0.93 | 0.66 | 0.84 | 0.80 | 0.96 | 0.87 |
| 30 | C8 | 0.83 | 0.93 | 1 | 0.78 | 1 | 0.89 |
| 33 | C9 | 0.83 | 1 | 0.71 | 1 | 1 | 0.89 |
| 36 | C10 | 0.80 | 0.79 | 0.61 | 0.93 | 0.9 | 0.81 |
| 39 | C11 | 0.81 | 0.77 | 0.60 | 0.83 | 1 | 0.97 |
| 42 | C12 | 0.80 | 0.73 | 0.58 | 0.73 | 0.95 | 0.85 |
| 45 | C13 | 0.815 | 0.49 | 0.46 | 0.65 | 0.88 | 0.76 |

**Table 6.** The table shows the relationship between molecular target interventions and the probabilities of autophagy and neuron death. The data includes the levels of seven different inputs, which are known to play a role in these processes. The table presents the resulting probability of autophagy and neuron death for each combination of input levels

| Inputs |  |  |  |  |  |  | Outputs |  |
| --- | --- | --- | --- | --- | --- | --- | --- | --- |
| 4EBP2 | TFEB | gsk3b | Wnt | PHLPP | RPS6KB1 | LRKK2 | Autophagy | Neuron death |
| 1 | – | – | – | – | – | – | 0.3983 | 0.9557 |
| – | 1 | – | – | – | – | – | 0.3691 | 0.9934 |
| – | – | 0 | – | – | – | – | 0.6918 | 0.9592 |
| 1 | 1 | – | – | – | – | – | 0.5055 | 0.8083 |
| – | – | 0 | 1 | – | – | – | 0.9928 | 0 |
| – | 1 | – | – | 0 | 0 | – | 0 | 0.0364 |
| 1 | 1 | 0 | – | 0 | 0 | 1 | 0.9926 | 0.0364 |

#### 3.3.4 FOXO3 activity

The FOXO3 activity pathway was also evaluated in this study due to its dysregulation in Parkinson’s disease (PD) (Pino et al. (2014)). The key element considered in this validation was the FOXO3 biomolecule, which plays important roles in autophagy, cell cycle progression, apoptosis, and stress resistance in PD (Pino et al. (2014); Cheng (2022); Fasano et al. (2019)).

The simulation scenario involved perturbing FOXO3 and observing the effects on autophagy and RNA-mediated biomolecules as activity levels using CellCollective. These results were confirmed through an exhaustive attractor search analysis. The results showed the following behaviors that are in line with published literature (table 4):

- The simulation results showed that FOXO3 activation increased autophagy in mitochondria (Fasano et al. (2019)).
- FOXO3 activation also activated different biomolecules involved in RNA-mediated mechanisms, including BECN1, GABARAPL1, MAP1LC3A, BNIP3, ATG12, and MUL1, which are known to be important regulators of autophagy (Hou et al. (2020)).

In order to validate the relevance of biomolecules for studying PD mechanisms, four pathways were analyzed, including one metabolic and three signaling pathways. The simulated behavior of these pathways was compared to the expected behavior based on published literature (table 4). While the behavior of the pathways was largely consistent with published literature, some discrepancies were observed in the Wnt/PI3K pathway, specifically in the simulated behavior of LRRK2. The results suggested that they are trustworthy indicators of biological processes, as they align with available data. Nonetheless, there was a mismatch in the Wnt-PI3K/AKT signaling pathway wre the model did not match published literature, leading to its revision.

Several factors could contribute to these discrepancies, such as differences in experimental conditions or protocols, the use of different LRRK2 models or cellular systems, the complexity of the autophagy pathway, and limited understanding of the exact mechanisms by which LRRK2 regulates the autophagy pathway. To address these discrepancies, corrective measures were taken, including modifying the Boolean function to better represent the interactions between LRRK2, ARFGAP1, and autophagy (Stafa et al. (2012)). These results suggest that the simulated behavior of the pathways is largely consistent with published literature.

## 4 DISCUSSION

In this study, Boolean modeling was used to examine the complexity of PD by simulating the dynamic interactions between various biomolecules. These models were used to test hypotheses about the role of certain biomolecules in PD progression, by simulating the effects of these biomolecules and comparing the results to experimental or observational data. Additionally, the models were used to investigate the impact of multiple perturbations on the disease and to identify patterns that may not be evident from experimental or observational data alone.

### 4.1 Constructing Boolean models from knowledge repositories

The Boolean models were automatically constructed using the CaSQ tool (Aghamiri et al. (2020)). One of the key benefits of using the CaSQ tool is its ability to apply specific rewriting rules to simplify the model and make it more manageable. Another advantage is its ability to translate diagrams into the SBML-qual format, which is a widely adopted standard in the systems biology community for representing qualitative dynamic systems. This format allows for the description of the model structure (biomolecules and interactions) and the mathematical equations describing their behavior over time, making it easy to share and compare models with other researchers.

The transformation of the diagrams into Simple Interaction Format (SIF) using the CaSQ tool was also beneficial, as it allowed for the use of various tools to analyze the diagrams. By using different tools to analyze the diagrams, researchers can gain a more comprehensive view of the disease, leading to an improved understanding of its underlying mechanisms.

The reduction of Process Description notation into Activity Flow notation and inference of logical functions from interactions, in combination with the use of SBML-qual format, resulted in the creation of models that are both accurate and computationally efficient. This makes the generated models suitable for studying large and complex systems, and allows for a deeper understanding of the system and more accurate predictions about its behavior. Therefore, the constructed Boolean models have the potential to be expanded to study larger and more complex systems.

It’s worth mentioning that using specific knowledge resources, such as PD map, has the advantage of providing a high quality of disease-specific knowledge. This is because the diagrams are focused on a specific disease and are created and reviewed by experts in the field. This expertise and specificity make the generated models more reliable.

### 4.2 Validation of the constructed models

The construction of models in compliance with systems biology standards enhances their compatibility with various tools and programs, leading to improved reproducibility and easier pipeline development. By adhering to these standards, researchers can ensure that their models are easily comprehended and integrated with other models and data sources in a consistent and dependable manner. Inadequate modeling details can lead to erroneous predictions. Thus, it is crucial to prioritize model quality during construction to minimize false positives during simulation. Verification is a critical aspect of the modeling process, as it confirms the reliability of the model and its ability to make accurate predictions and inform decision-making.

This study evaluated the BMs by demonstrating their ability to replicate experimentally validated studies from literature. The results, as shown in table 4, indicate that the simulated behavior of the models aligns with expected behavior under both original and perturbed conditions. The simulation of known perturbations confirms the models’ ability to recreate known pathological conditions, enhances understanding of these conditions, and enables the identification of a reliable set of biomolecules and translation rules. The selection of PD map diagrams for downstream modeling and verification is crucial in comprehending the biological mechanisms underlying PD and identifying potential targets for therapy development. These diagrams were chosen based on their relevance to PD phenotypes such as mitochondrial dysfunction, dopamine dysregulation, alpha-synuclein aggregation, neuroinflammation, and oxidative stress (Antony et al. (2013); MacMahon Copas et al. (2021)).

### 4.3 Structural and functional validation of the Boolean modelling approach

#### 4.3.1 Evaluation of the TCA cycle model

The TCA cycle model from the PD map was used to validate the Boolean modeling approach. The TCA cycle is a widely studied pathway with ample experimental data on enzyme activity and metabolite levels, making it possible to compare BM predictions with experimental data and assess the accuracy and reliability of the model. Additionally, dysregulation of the TCA cycle is linked to oxidative stress, inflammation, and cell death, which are all hallmarks of Parkinson’s disease (Stacpoole (2012); Kim et al. (2016); Sergi and Parayil Sankaran (2021)).

The structural and functional roles of key molecules in the TCA cycle BM were analyzed. This involved examining the involved enzymes and cofactors, as well as the reactions they catalyze and the intermediates they produce. The regulatory mechanisms controlling the activity of these molecules were also considered to predict their impact on the TCA cycle (Kafkia et al. (2022)). The effects of overexpression and knockout of the regulatory mechanisms controlling the activity of the TCA cycle enzymes were modeled in the TCA cycle BM. The results from the structural analysis were used to calculate the sensitivity of the TCA cycle to knockouts and overexpressions by simulating the impact of knockouts and overexpressions of specific biomolecules on the overall activity of the TCA cycle. Literature was reviewed to find experimental studies that perturbed TCA cycle molecules in model organisms, such as yeast or mice, to verify the results (Wongkittichote et al. (2019); Lee et al. (2011)).Selected perturbations in the Boolean model resulted in the following findings supported by the literature (table 4):

- Activation of acetyl CoA, NADH, and PDKs in silico increased the phosphorylation reaction, reducing the activity of the pyruvate dehydrogenase complex (PDC), which led to decreased levels of succinic semialdehyde and succinic acid (Stacpoole (2012); Sergi and Parayil Sankaran (2021)).
- Simulated KGDHC knockout predicted a deficiency in succinic acid and succinyl CoA and a downstream decrease in ATP and GTP levels (Shi et al. (2011); Gibson et al. (2003)).
- Simulated isocitrate dehydrogenase knockout resulted in decreased ATP production and inhibited the oxidative decarboxylation of isocitrate, with decreased levels of oxoglutarate observed in response to L-glutamate (Kim et al. (2016); Huergo and Dixon (2015)).
- Knockout of SIRT3 downregulated oxoglutaric acid, glutamate hydrogenase 1 (GLUD), and ATP levels, directly impacting mitochondrial function (Shen et al. (2020)).
- The simulation of the effect of alpha KGDH and GTP, GDP on ATP levels was validated using a probabilistic BM based on temporal-fluxomics data, which describes the oscillations of metabolites in the TCA. The simulated activity levels matched the measured concentrations in synchronized HeLa cells at various time points post-release from growth arrest (table 5).

The TCA cycle BM used in this study was capable of reproducing molecular activity that affects ATP levels and mitochondrial function. These results indicate that the TCA BM is a trustworthy tool for depicting the general picture of energy metabolism and can shed light on the mechanisms behind the oscillations of cellular metabolites. Nevertheless, this evaluation has some limitations, such as the possibility of missing relevant data in the literature search or not accurately reflecting all complex interactions within cells. Additionally, the context of evaluating the TCA cycle may differ based on the organism or tissue being studied. Despite these limitations, the primary goal of validating the TCA cycle BM is to ensure its accuracy in reflecting the underlying biological processes.

### 4.4 Modeling of the signaling pathways

#### 4.4.1 Dopamine transcription: The role of the NR4A2 gene

The NR4A2 gene plays a critical role in the regulation of dopamine and the growth and preservation of neurons. This gene is expressed in various tissues, including the brain, and has been implicated in several vital biological processes, such as neuron survival, mitochondrial biogenesis, and apoptosis (Chen et al. (2020)). When modeled using in silico methods, the knockout of the NR4A2 gene correctly predicts the decrease in BDNF production (Barneda-Zahonero et al. (2012)) and the alteration of expression of other genes involved in neurotransmitter metabolism and transport, including dopamine (Jankovic et al. (2005); Kadkhodaei et al. (2013); Jacobs et al. (2009)). Activation of the SIRT1 gene, which is also involved in neurotransmitter metabolism, leads to improved mitochondrial biogenesis, a process required for the formation of new mitochondria.

#### 4.4.2 Wnt-PI3K/AKT signalling: Implications for dopaminergic neurogenesis and autophagy

The Pi3K/AKT pathway and Wnt signaling are known to be critical for dopaminergic neurogenesis, as well as for crucial developmental processes and the aging process, which is a major risk factor PD (Marchetti et al. (2020); Long et al. (2021)). Both pathways share common downstream targets, suggesting the possibility of crosstalk and potential synergistic therapeutic approaches. For instance, both pathways can regulate the activity of GSK3B, a signaling protein, and are involved in autophagy, protein translation, and neuronal survival (Castelo-Branco et al. (2004); Hermida et al. (2017)). Numerous studies have validated the content and structure of these pathways. Of note, Pi3K/AKT pathway and Wnt signaling have been shown to play a significant role in PD pathogenesis. The Pi3K/AKT pathway has been implicated in modulating neuronal survival, while Wnt signaling has been implicated in both neuroprotection and neurodegeneration. Additionally, the interaction of the two pathways with alpha-synuclein, a protein that aggregates and forms Lewy bodies, a hallmark feature of PD, suggests a complex interplay between these pathways in PD pathogenesis. Therefore, a deeper understanding of the interaction between the Pi3K/AKT pathway and Wnt signaling in PD pathogenesis could provide novel insights into disease mechanisms and novel therapeutic strategies (Marchetti et al. (2020); Long et al. (2021)).

The following points outline key findings from modeling studies on the role of various molecular pathways in Parkinson’s disease (PD):

- The LRRK2 gene, particularly its G2019S mutant form, has been associated with an increased risk of developing PD (Madureira et al. (2020)). Overexpression of the LRRK2 G2019S mutant has been shown to enhance autophagy, a process involving the degradation and recycling of cellular biomolecules, through the inhibition of mTORC1/2 (Bravo-San Pedro et al. (2013); Poret and Guziolowski (2018); McCabe et al. (2020); Zhu et al. (2019)). Simulations of LRRK2 overexpression predicted the reactivation of mTORC1, which is consistent with proposed interactions between amino acid sensing, mTORC1 signaling, and autophagy (Rabanal-Ruiz et al. (2017)).
- TFEB, a transcription factor that regulates autophagy, has been shown to have complex effects on this process, activating autophagy while also activating protein synthesis inducers such as EIF4E and RPS6KB1 (Decressac et al. (2013)). Simulations of TFEB overexpression predicted a neuroprotective effect through increased autophagy activity, while joint activation of TFEB and protein synthesis inhibitor 4EBP2 led to a further increase in autophagy activity (Franco-Juárez et al. (2022); Zhuang et al. (2020); Decressac et al. (2013)). The inhibition of RPS6KB1 and PHLPP and activation of TFEB significantly decreased neuronal death (Decressac et al. (2013)).
- The Wnt signaling pathway and GSK3B have also been implicated in PD. Simulated overexpression of Wnt protein and inactivation of GSK3B were shown to increase autophagy (Awad et al. (2017); Moya et al. (2014); Momčilović et al. (2014)). Simulations of GSK3B inhibition and overexpression indicated its role in regulating neurogenesis, consistent with reported findings (Castelo-Branco et al. (2004); Toledo et al. (2017)). Modulating Wnt signaling and GSK3B together may hold potential as a neuroprotective treatment in early stages of PD progression (Awad et al. (2017); Moya et al. (2014); Momčilović et al. (2014); Castelo-Branco et al. (2004); Toledo et al. (2017)).

In conclusion, the validation study of the Wnt/PI3K pathway has confirmed existing literature findings and has also generated novel hypotheses regarding the combined modulation of its components. However, it is important to note that the relationship between the discussed biomolecules and neurogenesis and autophagy is complex and may depend on specific cell types, activity of other signaling pathways, or the stage of disease progression. Further studies are needed to fully elucidate the mechanisms underlying the interactions between these signaling pathways and their downstream targets, and to determine their potential as therapeutic targets in PD. Nonetheless, the findings presented in this study provide a promising avenue for further exploration and potential development of novel treatments for PD.

#### 4.4.3 FOXO3 activity: Impact on mitochondrial autophagy

The FOXO3 activity pathway has been identified as a critical mediator of mitochondrial homeostasis (Fasano et al. (2019)). The BM of FOXO3 activation have revealed that this pathway specifically promotes autophagy in mitochondria, through the activation of various biomolecules, including BECN1, GABARAPL1, MAP1LC3A, BNIP3, ATG12, and MUL1 (Hou et al. (2020)). These biomolecules have been established as key regulators of autophagy. Thus, the activation of these biomolecules in response to FOXO3 activation highlights a close interconnection between RNA-mediated pathways and FOXO3 activity in the regulation of autophagy. These findings have the potential to improve our understanding of the complex mechanisms involved in the regulation of autophagy and its role in maintaining cellular health. Further investigations are required to clarify the extent and specifics of this relationship, which could enhance the development of targeted therapeutic strategies for disorders related to mitochondrial dysfunction and autophagy.

### 4.5 Applications and limitations of results in translational research

Translational medicine aims to bridge the gap between basic research and clinical practice by applying scientific findings to medical practice. One way to achieve this goal is through the use of modeling and simulation techniques. These techniques use existing knowledge from bench experiments and disease-relevant omics datasets to develop new hypotheses about the disease and propose improved therapies and diagnostics. The Boolean modeling approach, which is based on systems biology and systems medicine, is particularly useful because it allows for iterative improvements in understanding through a continuous cycle of data-driven modeling and model-driven experimentation. The Boolean modeling approach has the potential to generate therapy-related hypotheses by designing perturbation experiments to compare model attractors to the disease signature. This allows the identification of the basins of attraction that could alleviate the pathological signature of the disease and suggest the best combinations of targets to achieve a healthy state.

This study offers valuable insights into the potential use of a standardized SBML-qual format for Parkinson’s disease pathway models. Through the use of the CaSQ translation tool, the authors were able to generate models that can inform future research efforts. Moreover, the study highlights the importance of accurate and relevant data input, appropriate assumptions and algorithms, and further integration with omics data for improving the accuracy and relevance of modeling and simulation approaches in understanding disease mechanisms and proposing targeted therapies. We also emphasize the need for disease-specific datasets to accurately parametrize models and better understand individual dysregulations in different disease subtypes. This personalized approach has the potential to lead to the development of more effective treatments tailored to the unique characteristics of each disease subtype.

While the study acknowledges certain limitations, such as the high-level representation of observable disease effects and variations in disease mechanism dynamics between patients, it highlights the potential for continued improvement in modeling and simulation techniques to advance translational medicine and improve disease diagnosis and therapy. By addressing these limitations and incorporating more precise and individualized data, modeling and simulation approaches can provide a powerful tool for advancing our understanding of disease progression and the development of targeted therapeutic approaches.

## 5 CONCLUSION

The findings of this study demonstrate that Boolean modeling is a powerful and promising tool for understanding the complexity of Parkinson’s disease. By modeling the dynamic interactions between various biomolecules, researchers can gain insight into the underlying mechanisms of the disease and propose potential therapeutic strategies.

Boolean models allow us to test hypotheses about the role of particular biomolecules in the development of PD. The models can also be used to explore the impact of multiple perturbations on the disease and identify patterns that may not be apparent from experimental or observational data alone

Despite its potential, several challenges remain in the use of Boolean models, including improving the accuracy and predictive power of these models. Model refinement, data integration, and parameter optimization can be applied to address these challenges. Model stratification is also a promising future direction for the field, which has the potential to increase the accuracy of the models and allow for more precise treatments.

Improving the interoperability, annotations, and reproducibility of Boolean models can enhance their accuracy. Standardized annotations can provide more information about the model, making it easier to understand and use. Improved interoperability can help to integrate different models and datasets, leading to more accurate predictions. Reproducibility can help to validate the results of the model and increase confidence in its predictive power. By improving these parameters, we can increase the accuracy and reliability of Boolean models, ultimately leading to better predictions and insights into the underlying mechanisms of complex diseases.

## AUTHOR CONTRIBUTIONS

Ahmed Abdelmonem Hemedan: Investigation, Writing – original draft. Reinhard Schneider: Supervision, Writing – review & editing. Marek Ostaszewski: Conceptualization, review & editing, Supervision.

## DATA AVAILABILITY STATEMENT

The data and scripts used to generate the results for this study are available in the LCSB GitLab repository (https://gitlab.lcsb.uni.lu/lcsb-biocore/boolean-modelling-of-pd-paper). The repository includes the fluxomics data used in the study, which was originally published by Ahn et al. in 2017 (Ahn et al. (2017)), and was used for the modeling and validation of the TCA cycle. In addition, boolean models generated from the PD map are also included in the repository. Access to this data can also be found on the LCSB GitLab repository.

