## Supplementary figures and tables for "Applications of Boolean modeling to study the dynamics of a complex disease and therapeutics responses"

### Supplementary Material

#### 1 SUPPLEMENTARY TABLES AND FIGURES

#### 2 SELECTED DIAGRAMS FROM PD MAP

We include the following pathway diagrams from PD map:

- Dopamine transcription pathway (fig. S1)
- FOXO3 activity pathway (fig. S2)
- PI3KAKT signalling pathway (fig. S3)
- mTOR pathway (fig. S4)
- PRKN signalling (fig. S5)
- TCA cycle (fig. S6)

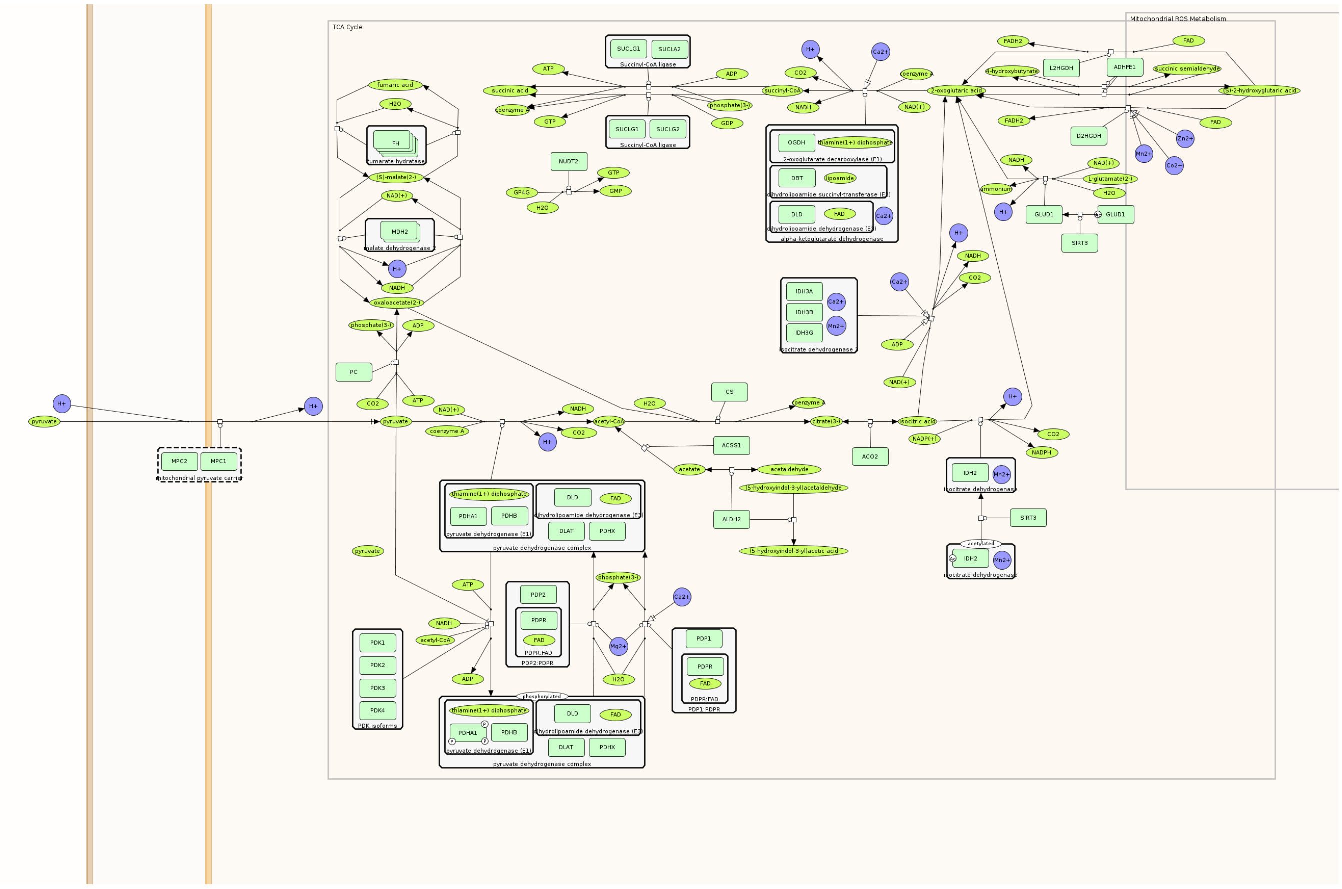

Figure S1: Dopamine transcription pathway

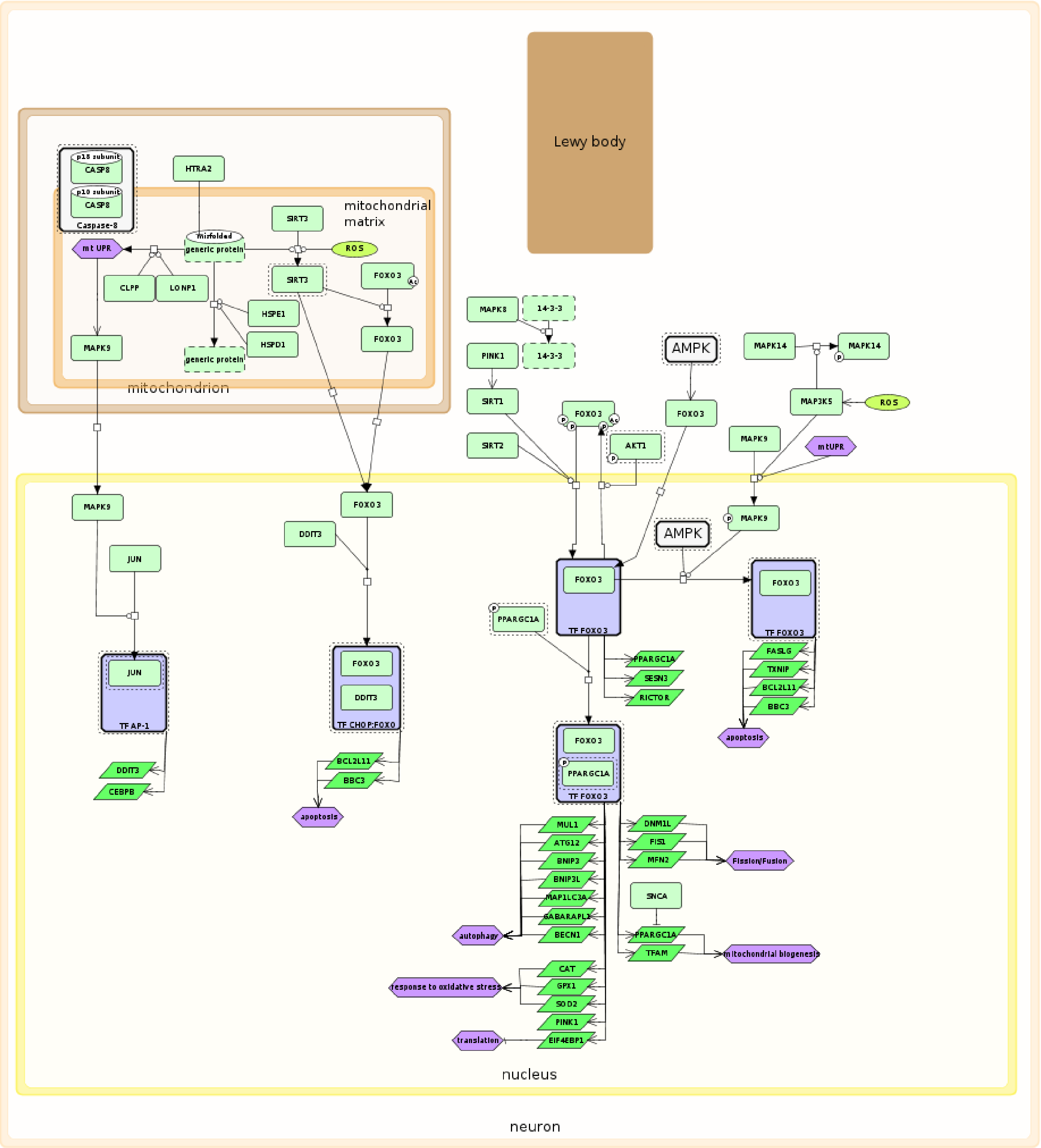


Figure S2: FOXO3 Activity pathway

#### 3 THE SIMULATION GRAPHS FROM BOOLEAN SIMULATIONS

This section details the simulation graphs that resulted from Boolean models and Probabilistic BMs simulation. All simulation graphs are also available in the gitlab repository .

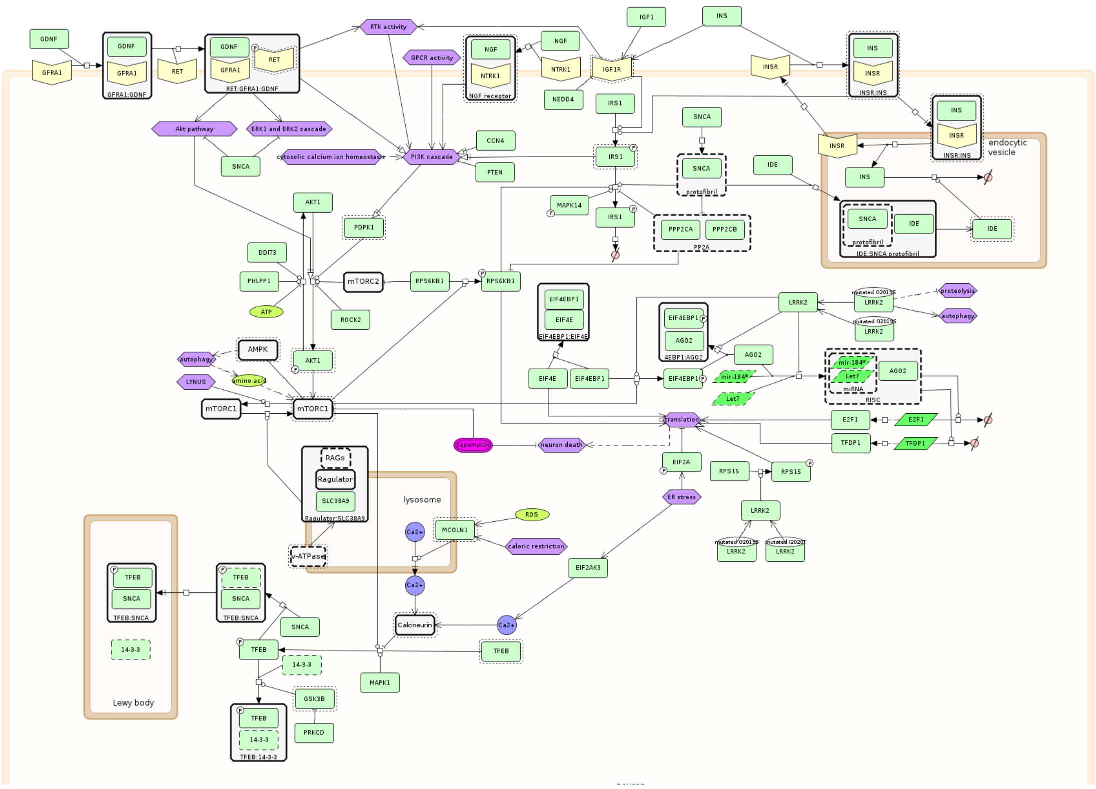


Figure S3: PI3KAKT signalling pathway

The examples from simulation graphs from CellCollective platform are shown in figs. S7 to S10. Interactive demo and guidance on how to construct and simulate the models is available at `https://cellcollective.org/`.

#### 4 THE SENSITIVITY AND STRUCTURAL ANALYSIS

Distances computed between the original and perturbed attractors are summarized in table S1 for knockouts and table S2 for overexpressions.

Identifying high betweenness centrality molecules with low sensitivity is important as it suggests the presence of compensatory paths. This information can be used to develop targeted interventions to disrupt the function of the pathway in pathological conditions. The results of our study highlight potential intervention points in different pathways, such as Wnt, ppargc1a, mTOR, Foxo3, and dopamine.

For instance, in the Wnt Pi3kAKT pathway, molecules such as GRM3, TGFB1 rna, IGF1, and INS compartment can compensate for the absence of PDPK1, RPS6KB1 phosphorylated, and IRS1 phosphorylated. In the PPARGC1 pathway, different subunits of the complex itself, various mitochondrial genes, and transcription factors such as TF NRF1 complex can compensate for the loss of Complex IV. In the mTOR pathway, compensatory molecules include other upstream signals, SIRT1 and PPARGC1 for AMPK complex neuron, other upstream signals for TSC1 TSC2 complex neuron, and other molecules involved in autophagy and lysosome function for SEN2. Similarly, in the FOXO3 pathway, TF CHOP FOXO complex and other pro-apoptotic molecules such as BCL2L11 rna and FASLG rna can compensate for the absence of BCL2L11 rna and BBC3 rna, and other forms of FOXO3 or other transcription factors for FOXO3 acetylated phosphorylated and MAPK9 phosphorylated. Finally, compensatory molecules for

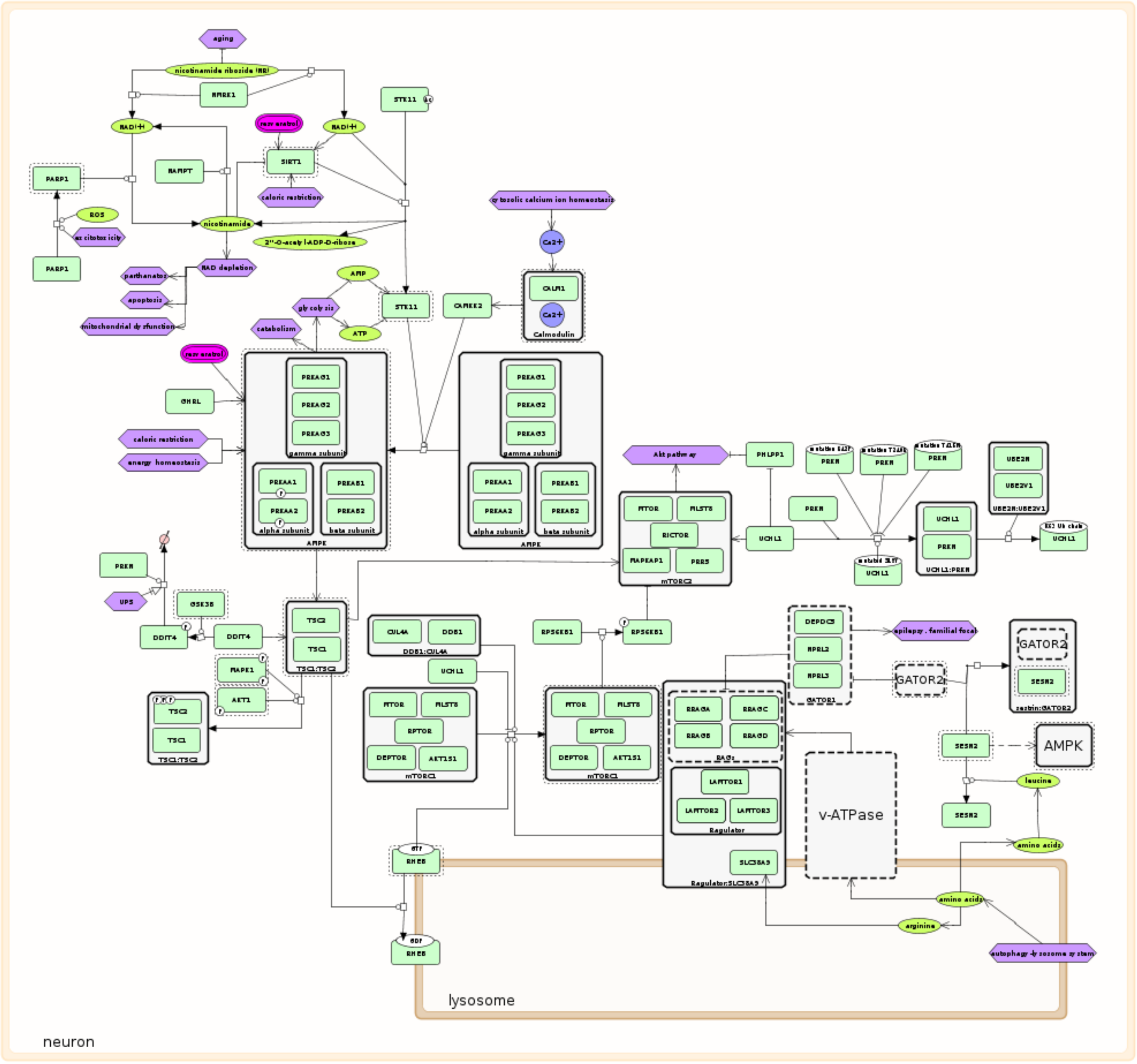


Figure S4: mTOR pathway

the dopamine transcription pathway involve other molecules in dopamine metabolism, neuron survival, and retinoic acid synthesis phenotypes such as SLC18A2 rna, GDNF rna, and TF PITX3 complex for BDNF rna, SLC18A2 rna, SLC6A3 rna, DRD2 rna, ALDH1A1 rna, and TF PITX3 complex for TH rna, and retinoic acid and TF NR4A2 complex for RXRA.

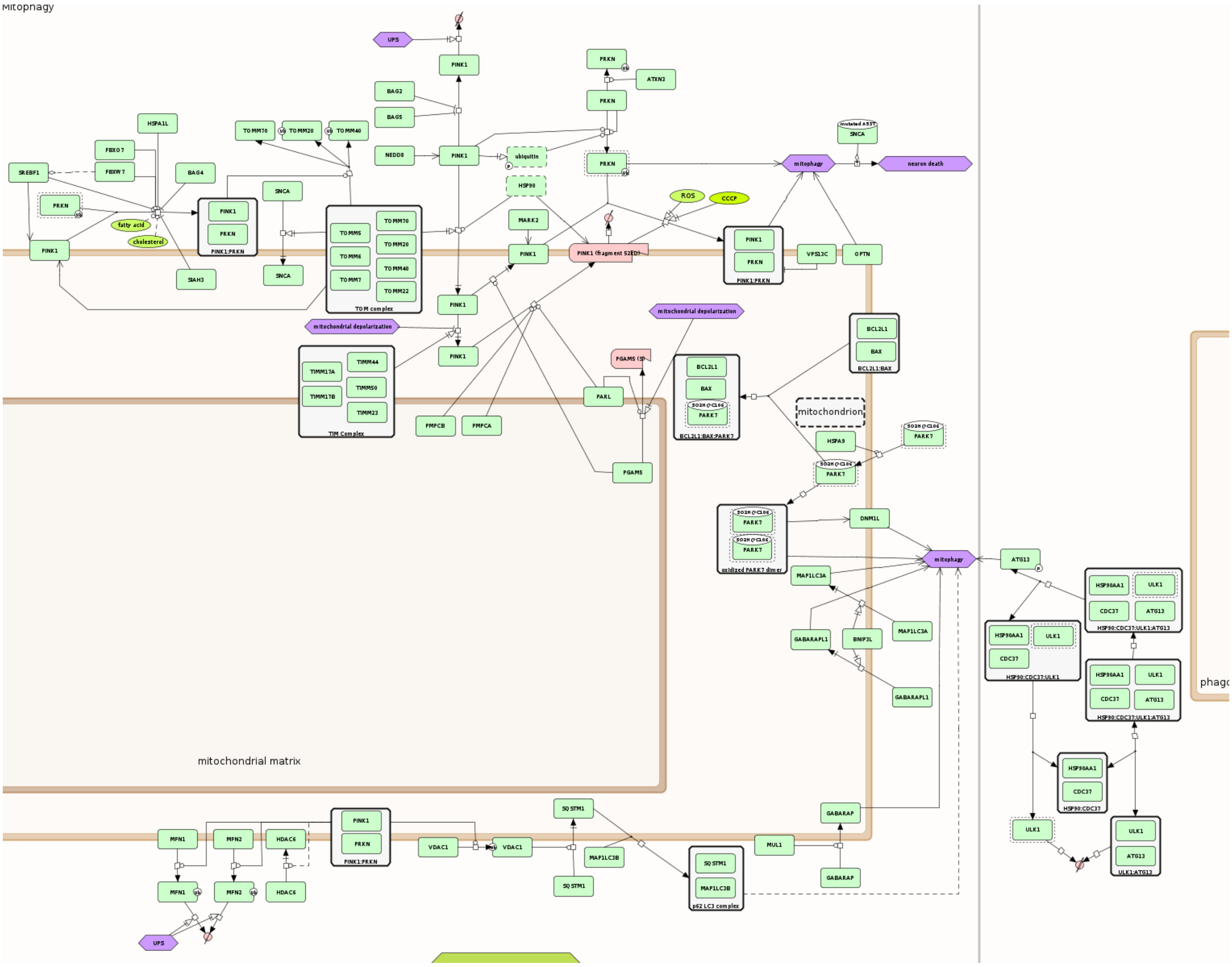

Figure S5: PRKN signalling pathway

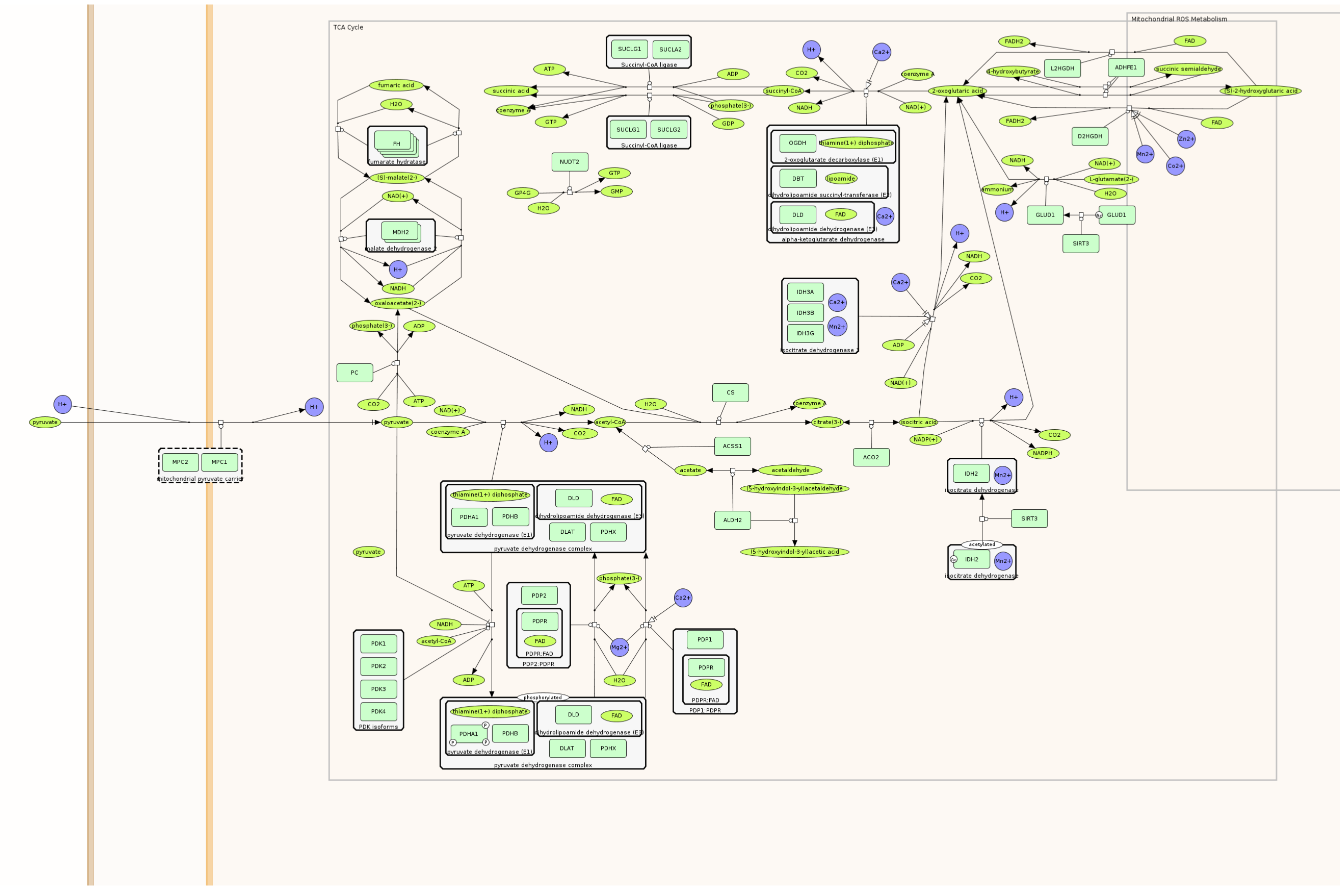

Figure S6: TCA cycle

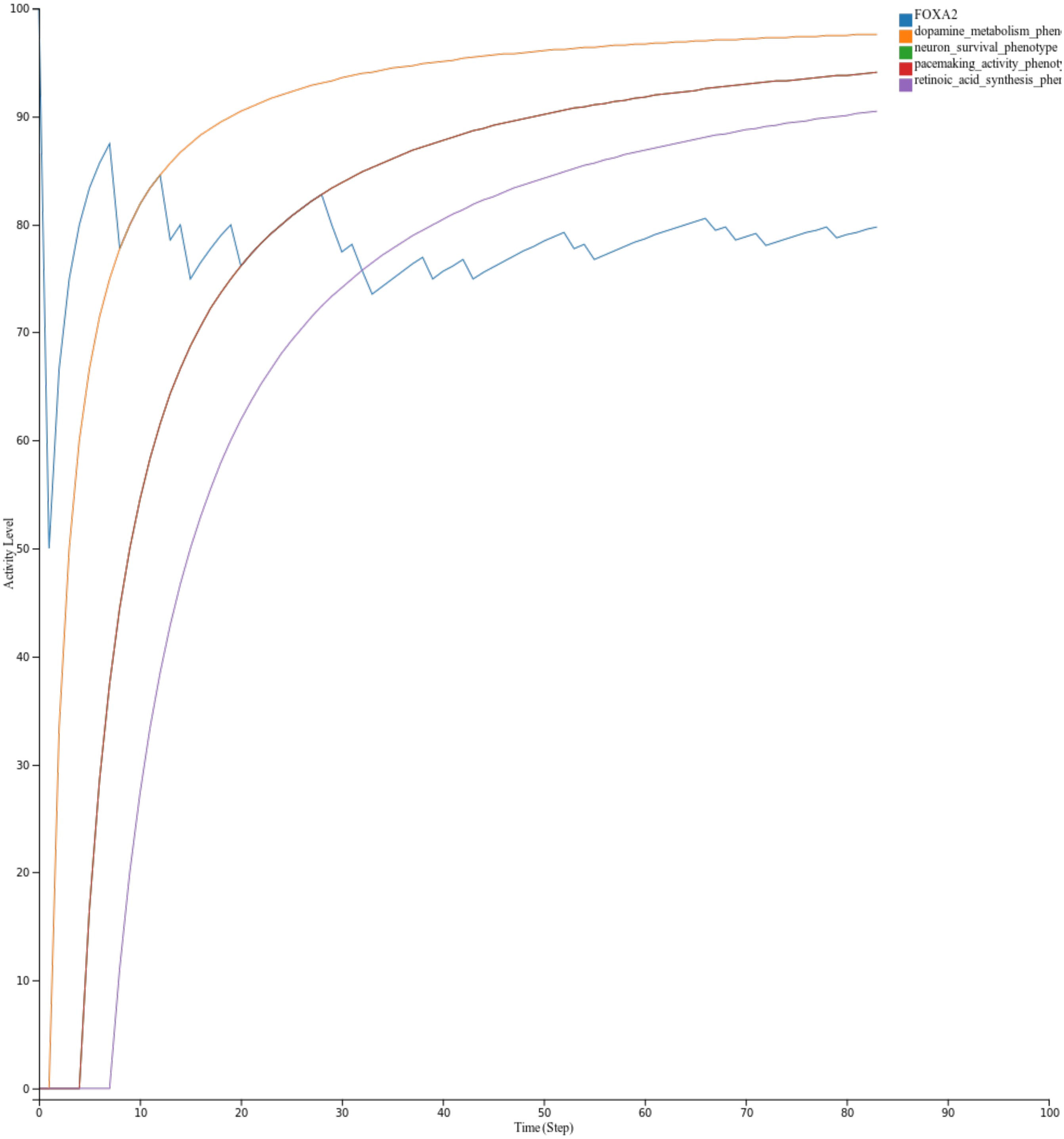


Figure S7: Dopamine transcription simulation

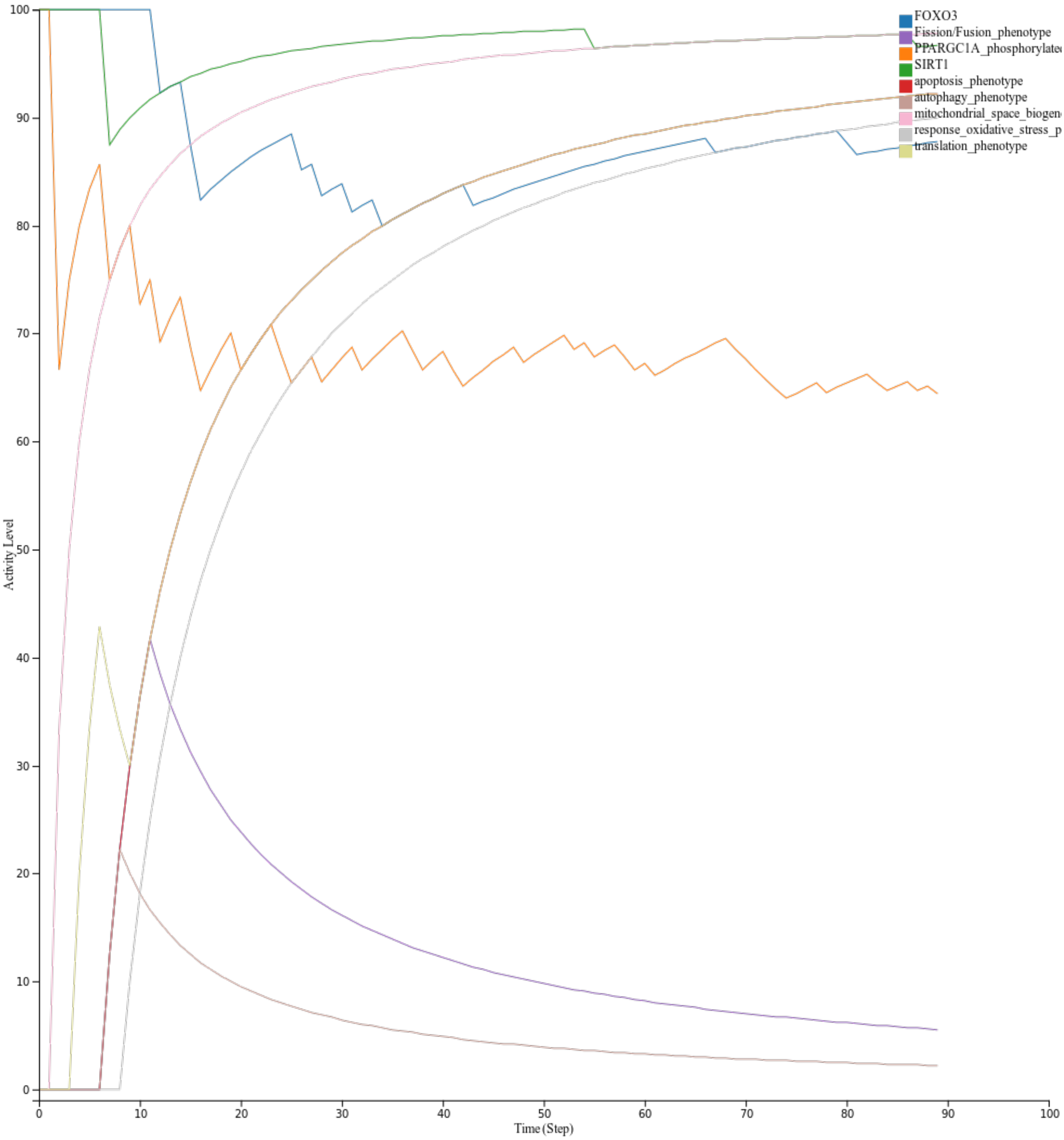


Figure S8: FOXO3 simulation

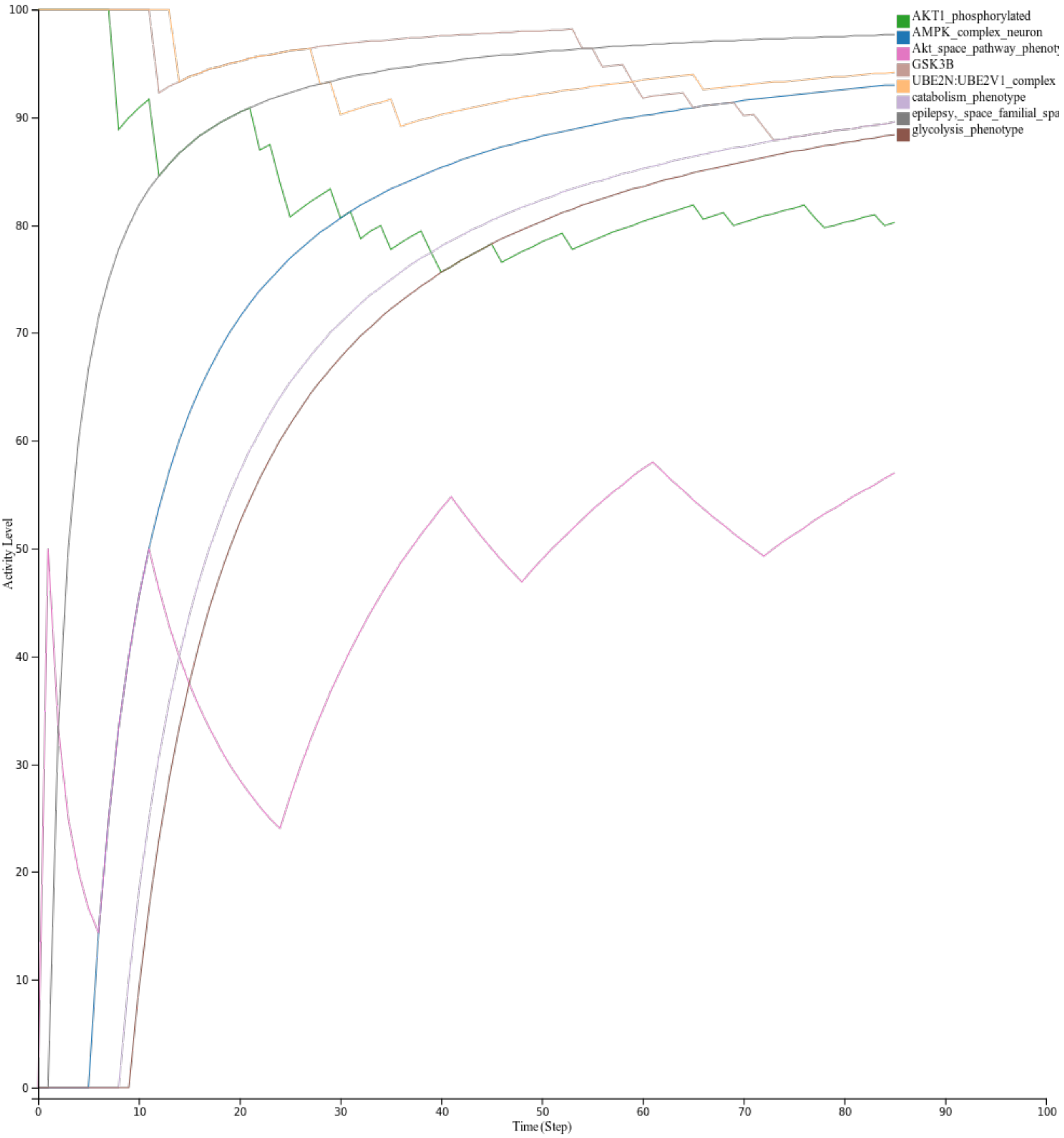


Figure S9: mTOR simulation

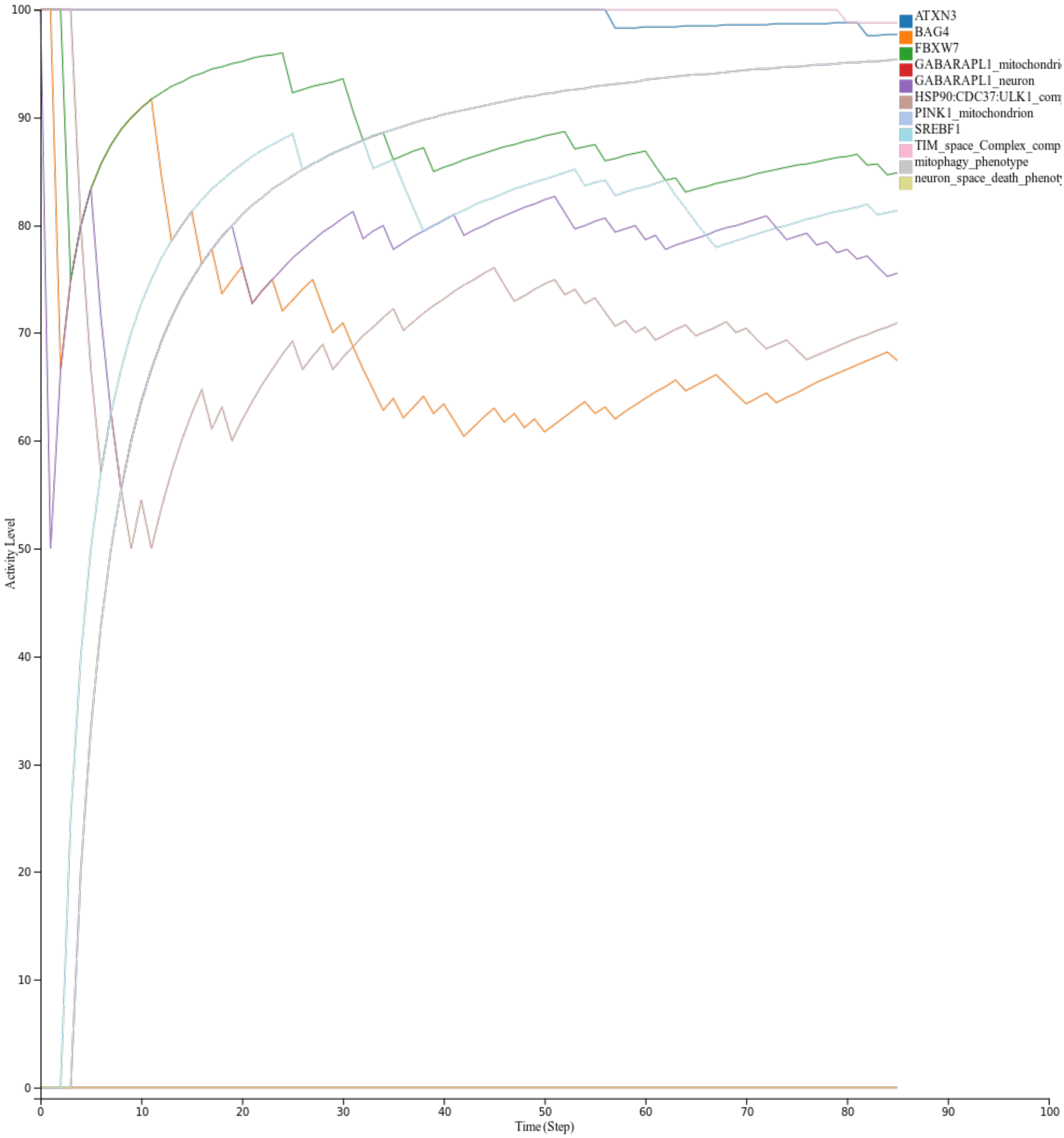


Figure S10: PRKN simulation

| Pathway | Group ID | Identity-based distance | Similarity-based distance |
|---|---|---|---|
| Pi3k/akt | RPS6KB1 | 0.582524272 | 0.012230182 |
| | PHLPP1 | 0.582524272 | 0.007517202 |
| | WNT1 | 0.563106796 | 0.013361297 |
| | WNT3 | 0.553398058 | 0.008483363 |
| | PRKN | 0.553398058 | 0.005372797 |
| TCA cycle | PDP2:PDPR_complex | 0.630769231 | 0.010729783 |
| | alpha-ketoglutaratedehydrogenase_complex | 0.630769231 | 0.009704142 |
| | PDP1:PDPR_complex | 0.6 | 0.01025641 |
| | SIRT3 | 0.6 | 0.037790927 |
| | isocitratedehydrogenase_complex | 0.6 | 0.012544379 |
| | GLUD1 | 0.6 | 0.01530572 |
| PRKN | SREBF1 | 0.648148148 | 0.012003 |
| | FBXW7 | 0.648148148 | 0.024005 |
| | OPTN | 0.648148148 | 0.012003 |
| | HSP90 | 0.611111111 | 0.012003 |
| PPARGC1A | IDH3G_rna | 0.582089552 | 0.008539393 |
| | TF_YY1_complex | 0.582089552 | 0.060741071 |
| | IDH3A_rna | 0.582089552 | 0.008539393 |
| | NDUFS8_rna | 0.582089552 | 0.008539393 |
| | ATP5MC1_rna | 0.582089552 | 0.008539393 |
| mTOR | TSC1:TSC2_complex_neuron | 0.603174603 | 0.0095742 |
| | SESN2 | 0.603174603 | 0.0095742 |
| | NAMPT | 0.587301587 | 0.009322247 |
| | ROS | 0.571428571 | 0.009070295 |
| | Akt | 0.571428571 | 0.009070295 |
| Foxo3 | MAP3K5 | 0.61971831 | 0.008728427 |
| | EIF4EBP1_rna | 0.605633803 | 0.008530054 |
| | ATG12_rna | 0.605633803 | 0.008530054 |
| | BECN1_rna | 0.605633803 | 0.008530054 |
| | BBC3_rna | 0.591549296 | 0.00833168 |
| | JUN | 0.577464789 | 0.008133307 |
| | SIRT1 | 0.577464789 | 0.008133307 |
| Dopamine transcription | EN1 | 0.852941176 | 0.013327206 |
| | LRRK2 | 0.573529412 | 0.046243107 |
| | FOXA2 | 0.573529412 | 0.008961397 |
| | SNCA | 0.558823529 | 0.022575827 |
| | SFPQ | 0.529411765 | 0.008272059 |
| | PIN1 | 0.514705882 | 0.008042279 |
| | RXRA | 0.5 | 0.0078125 |

Table S1: Examples shows the significant distances between the original and perturbed attractors (Knockouts)

| Pathway | Group ID | Identity-based distance | Similarity-based distance |
|---|---|---|---|
| Pi3k/akt | CTNNB1_phosphorylated | 1 | 0.015081535 |
| | EIF4EBP1_phosphorylated | 1 | 0.017862192 |
| | IRS1_phosphorylated | 1 | 0.009708738 |
| | CTNNB1_ubiquitinated_phosphorylated | 1 | 0.009708738 |
| | PDPK1 | 1 | 0.014209633 |
| | RPS6KB1_phosphorylated | 1 | 0.010085776 |
| | AKT1_phosphorylated | 0.990291262 | 0.018828353 |
| | TFEB_complex | 0.980582524 | 0.009685173 |
| TCA cycle | 2-oxoglutaricacid | 1 | 0.035108481 |
| | hydroxyglutaric_acid | 1 | 0.033609467 |
| | succinic_semialdehyde | 1 | 0.025325444 |
| | acetyl-CoA | 1 | 0.016094675 |
| | oxaloacetate_2 | 1 | 0.015384615 |
| | succinyl-CoA | 1 | 0.015384615 |
| | succinic_acid | 1 | 0.015384615 |
| | GMP | 0.984615385 | 0.014674556 |
| | GTP | 0.984615385 | 0.014674556 |
| PRKN | PRKN_ubiquitinated | 0.981481481 | 0.021604938 |
| | PINK1_neuron | 0.944444444 | 0.043552812 |
| | ubiquitin_phosphorylated | 0.944444444 | 0.017489712 |
| | PGAM5__S_ | 0.907407407 | 0.016803841 |
| | PINK1_mitochondrion | 0.888888889 | 0.017489712 |
| | PINK1 | 0.87037037 | 0.01611797 |
| PPARGC1A | COX5A_rna | 0.985074627 | 0.014702606 |
| | COX7A2_rna | 0.985074627 | 0.014702606 |
| | COX5B_rna | 0.985074627 | 0.014702606 |
| | CYCS_rna | 0.985074627 | 0.015296651 |
| | complex_IV_complex | 0.985074627 | 0.014702606 |
| | SDHB_rna | 0.985074627 | 0.014702606 |
| mTOR | CAMKK2 | 0.634920635 | 0.010078105 |
| | MAPK1_phosphorylated_phosphorylated | 0.603174603 | 0.0095742 |
| | AMPK_complex_neuron | 0.571428571 | 0.009070295 |
| | DDB1:CUL4A_complex | 0.571428571 | 0.009070295 |
| | PRKN_neuron | 0.555555556 | 0.008818342 |
| FOXO3 | FASLG_rna | 0.661971831 | 0.009323547 |
| | MAPK9_phosphorylated | 0.647887324 | 0.009125174 |
| | FOXO3_neuron | 0.591549296 | 0.00833168 |
| | PPARGC1A_rna | 0.591549296 | 0.00833168 |
| | FOXO3_acetylated_phosphorylated | 0.577464789 | 0.008133307 |
| | BCL2L11_rna | 0.577464789 | 0.008133307 |
| | FIS1_rna | 0.577464789 | 0.008133307 |
| Dopamine transcription | ALDH1A1_rna | 1 | 0.03125 |
| | DRD2_rna | 1 | 0.015625 |
| | TH_rna | 1 | 0.015625 |
| | BDNF_rna | 0.985294118 | 0.015395221 |
| | DDC_rna | 0.970588235 | 0.014820772 |
| | SLC18A2_rna | 0.941176471 | 0.014705882 |
| | SLC6A3_rna | 0.941176471 | 0.014705882 |
| | PITX3 | 0.911764706 | 0.026711857 |
| | TF_NR4A2_complex | 0.897058824 | 0.242704504 |

Table S2: Examples shows the significant distances between the original and perturbed attractors (overexpressions)

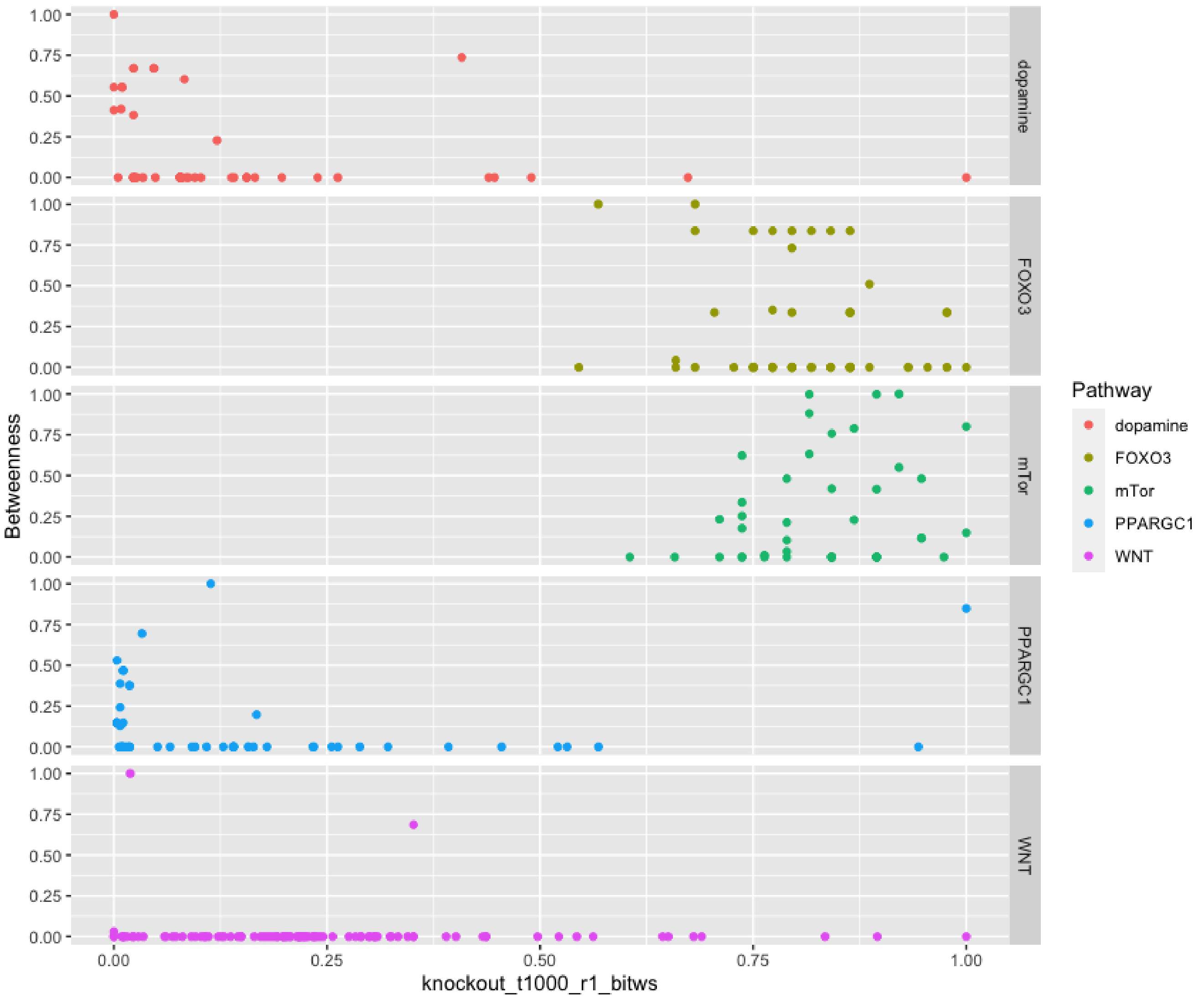


Figure S11: The figure represents molecules with high betweenness centrality and low knockout sensitivity in multiple pathways.

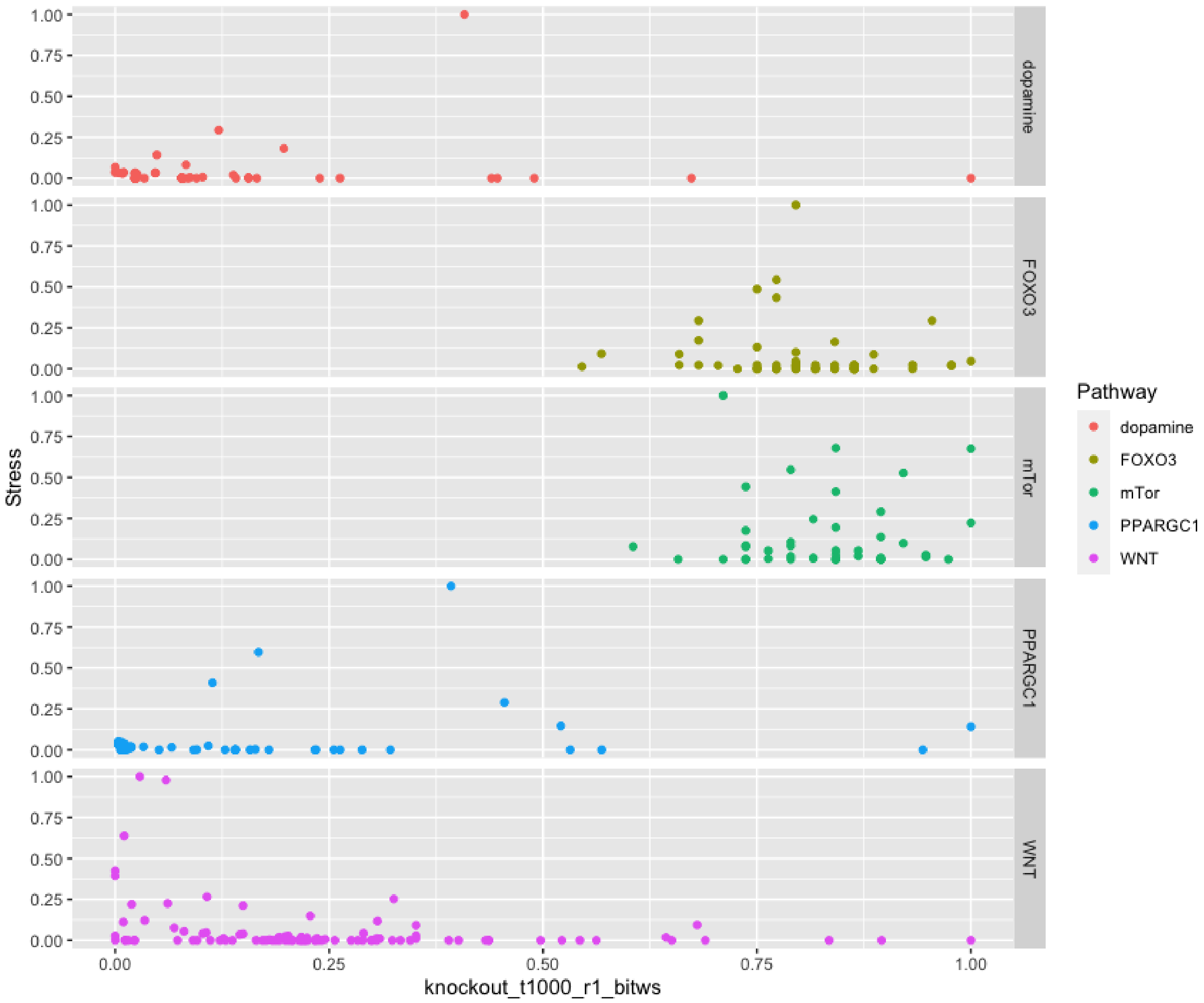


Figure S12: The figure represents the molecules with high stress centrality and low knockout sensitivities in multiple pathways suggests the presence of compensatory paths.

| Pathway | Node ID | Degree | | | Betweenness | Stress |
|---|---|---|---|---|---|---|
| | | Total | In | Out | | |
| Dopamine transcription | TF NR4A2 complex | 29 | 8 | 21 | 330 | 463 |
| | PITX3 | 3 | 2 | 1 | 102 | 136 |
| | TF PITX3 complex | 13 | 2 | 11 | 66 | 84 |
| | MIR133B rna | 2 | 1 | 1 | 54 | 66 |
| | RXRA | 2 | 1 | 1 | 27 | 38 |
| FOXO3 activity | TFFOXO3 complex nucleus | 36 | 9 | 27 | 3673 | 9084 |
| | TFCHOP:FOXO complex | 4 | 2 | 2 | 1763 | 4932 |
| | FOXO3 nucleus | 3 | 2 | 1 | 1600 | 4416 |
| | SIRT3 | 6 | 3 | 3 | 1482 | 3946 |
| | BBC3 rna | 3 | 2 | 1 | 896 | 2670 |
| mTOR | AMPK complex neuron | 10 | 7 | 3 | 2277 | 11064 |
| | TSC1:TSC2 complex neuron | 8 | 5 | 3 | 1465 | 7476 |
| | STK11 | 6 | 5 | 1 | 1018 | 6050 |
| | SESN2 | 5 | 2 | 3 | 788 | 2464 |
| | nicotinamide | 7 | 4 | 3 | 745 | 7518 |
| PPARGC1A | PPARGC1A phosphorylated | 13 | 4 | 9 | 322 | 650 |
| | TF NRF1 complex | 24 | 2 | 22 | 174 | 388 |
| | PPARGC1A(AC-Ph) | 4 | 3 | 1 | 94 | 188 |
| | TF NRF2 complex | 16 | 2 | 14 | 88 | 266 |
| | TF YY1 complex | 11 | 3 | 8 | 75 | 92 |
| TCA cycle | 2-oxoglutaricacid | 25 | 19 | 6 | 451 | 761 |
| | S-malate | 12 | 7 | 5 | 195 | 490 |
| | NADH | 19 | 15 | 4 | 188 | 539 |
| | ADP | 15 | 8 | 7 | 178 | 393 |
| | acetyl-CoA | 10 | 6 | 4 | 152 | 384 |
| | mTORC1 complex neuron | 11 | 7 | 4 | 463 | 511 |
| | AKT1 phosphorylated | 7 | 5 | 2 | 440 | 500 |
| Wnt-PI3K/AKT | PI3K | 9 | 8 | 1 | 297 | 369 |
| | PDPK1 | 2 | 1 | 1 | 260 | 326 |
| | RPS6KB1 phosphorylated | 5 | 3 | 2 | 189 | 201 |

Table S3: Common top five topological metrics in BMs and their source diagrams. The table summarises topological properties of nodes in the selected pathways, and “Node ID” indicate the specific nodes, “Degree” indicates a specific type of a node degree of in the BMs (total/incoming/outgoing connections), “Betweenness” describes node betweenness, while “Stress” describes the number of shortest paths that pass through a node

#### 5 ATTRACTOR IDENTIFICATION

The performance of asynchronous and synchronous simulations was evaluated in selected models to gain a comprehensive understanding of their characteristics and to determine their performance. The attractor analysis results indicated that the state trajectories converge to either fixed or cyclic attractors, dependent on the synchronous and asynchronous updating schemes. The comparison of four algorithms (HyTarjan, Heuristic, Decomp, and SAT) (**??** in terms of their calculation speed in pathways is summarized in table S4. A comparison of four different algorithms, namely HyTarjan, Heuristic, Decomp, and SAT, in terms of their calculation speed in pathways is presented in table S4. The algorithms and the data used for this comparison were obtained from the studies by (**??**, which evaluated the performance of these algorithms in Boolean network modeling. In our study, the purpose of this comparison is to identify which algorithm performs better in terms of speed when applied to pathway analysis of the Parkinson's disease. The results of this study may provide valuable insights into the selection of appropriate algorithms for this type of analysis.

The speed of the algorithms was compared. The SAT algorithm demonstrated improved time to find the attractors compared to the Decomp algorithm, with the exception of the TCA cycle. The SAT algorithm achieved a substantial 87.32% improvement in the ER stress signaling pathway, significantly reducing the time required to reach the attractor.

| | | | Time (seconds) | | | |
|---|---|---|---|---|---|---|
| | | | Asynchronous | | Synchronous | |
| Pathway | Edges | Targets | HyTarjan | Heuristic | Decomp | SAT |
| PGC1 alpha | 109 | PPARGC1A | 3547 | 1789 | 173 | 96 |
| | | SIRT1 | 2587 | 1471 | 169 | 74 |
| Dopamine transcription | 167 | NR4A2 | 2981 | 1460 | 147 | 54 |
| Wnt/PI3K-AKT | 391 | Wnt/PI3K | 1135 | 3961 | 403 | 256 |
| ER stress signaling | 53 | DDIT3 | 971 | 1855 | 67 | 11 |
| | | AKDHC | 2066 | 1123 | 84 | 110 |
| TCA cycle | 137 | Oxoglutarate | 2122 | 1140 | 84 | 110 |
| | | IDH | 2153 | 1151 | 84 | 110 |
| | | SIRT3 | 2130 | 1140 | 84 | 110 |

Table S4: Attractor reachability speed in different algorithms. The table shows the duration of attracor reachabiliy for asynchronous and synchronous systems in the selected pathways, using the methods HyTarjan, Heuristic, Decomp, and SAT. The scales includes the node numbers, and Targets indicate the perturbed molecules

The landscape of the network state transitions along with attractor cycles were identified. The returned transition network object has same structures with the normal network object. The transition network is written as a SIF file. The SIF file could be loaded to Cytoscape with the following the steps in fig. S13.

The result is shown in fig. S14.

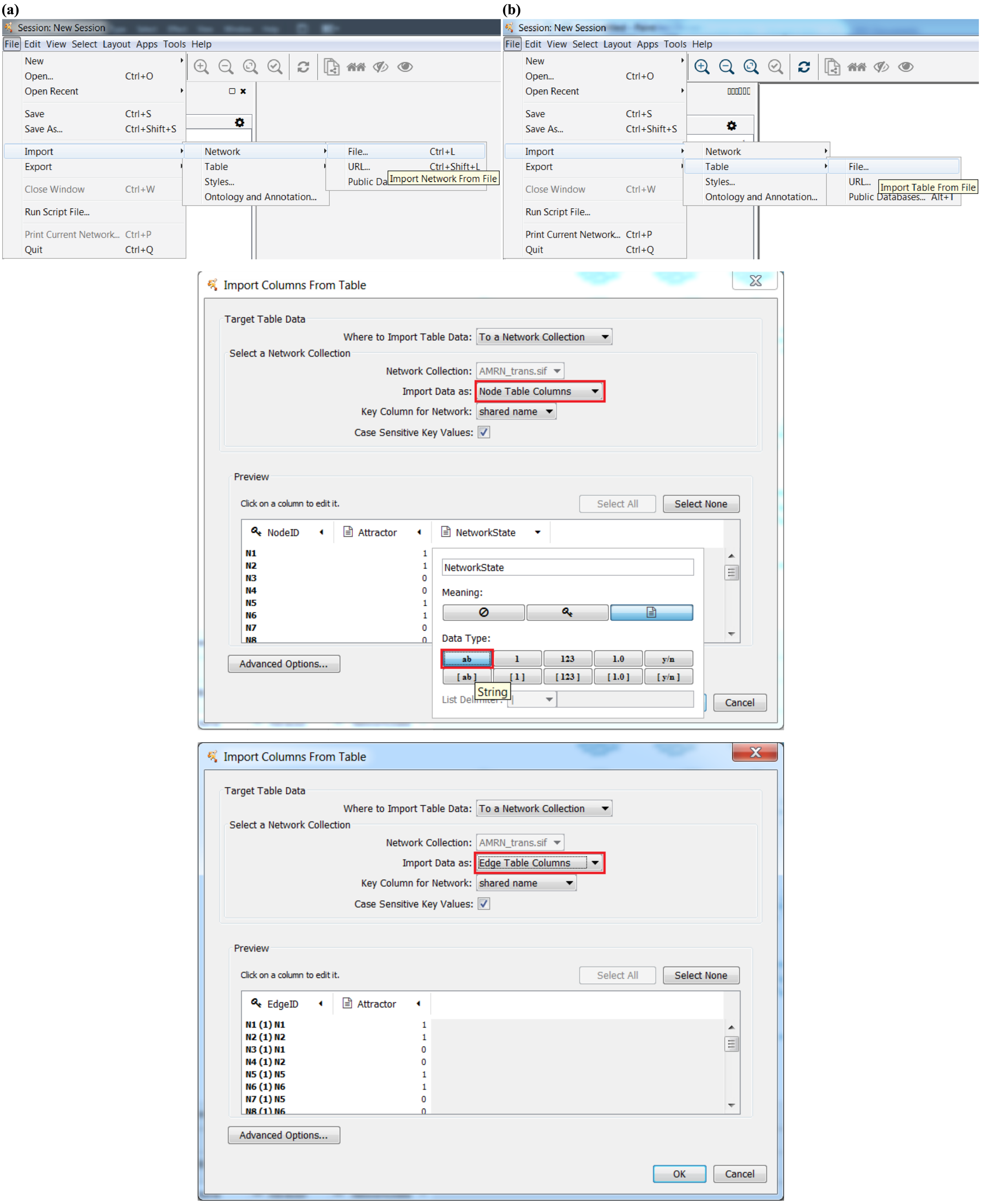


Figure S13: Steps for loading a SIF fyle in Cytoscape (top to bottom: opening the import dialogs, importing the nodes, importing the edges).

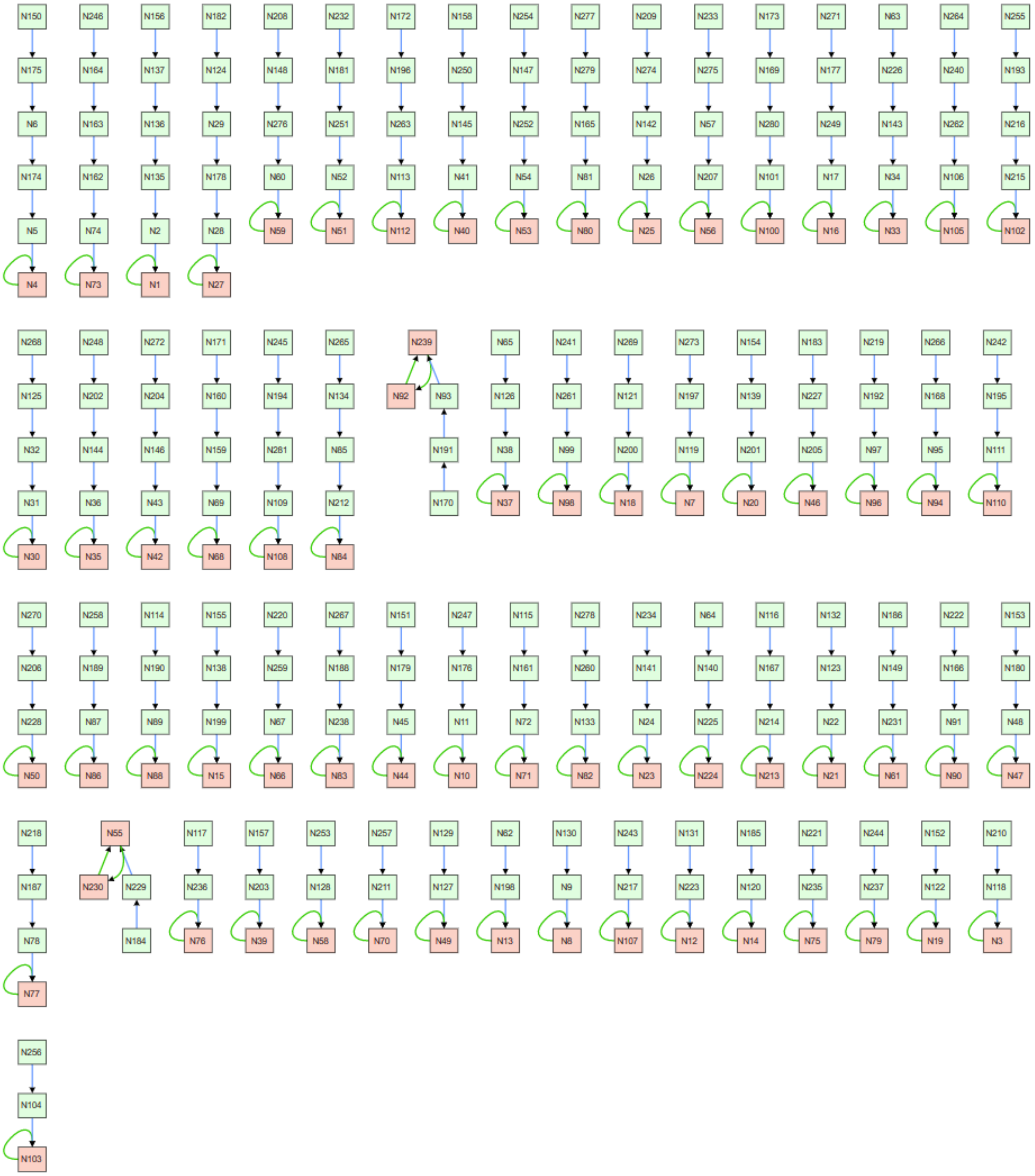

Figure S14: PPARGC1A pathway attractor
